# Mechanism of phosphoinositide regulation of lysosomal pH via inhibition of CLC-7

**DOI:** 10.1101/2025.10.01.679551

**Authors:** Jacob K. Hilton, Yifei Lin, Eric Sefah, Justin C. Deme, Joanne L. Parker, Matthew J. Langton, Michael Grabe, Susan Lea, Simon Newstead, Joseph A. Mindell

## Abstract

Lysosomes process cellular waste and coordinate responses to metabolic challenge. Central to lysosomal homeostasis are phosphoinositide lipids, key signaling molecules which establish organelle identity, regulate membrane dynamics and are tightly linked to the pathophysiology and therapy of lysosomal storage disorders, neurodegeneration, and cancer. Phosphatidylinositol 3,5- bisphosphate (PI(3,5)P2) interacts with multiple lysosomal membrane proteins and plays a critical role in regulating lysosomal pH by directly inhibiting the chloride/proton antiporter ClC-7, though the molecular mechanism of this inhibition remains unclear. Here, using a combination of functional, structural, and computational analysis, we demonstrate that PI(3,5)P2 binding dramatically remodels the structure of ClC-7 by inducing close association between cytosolic and transmembrane domains. Disease-causing mutations show increased transport activity through loss of PI(3,5)P2 binding and subsequent inhibition. Conversely, ClC-7 activation is correlated with dissociation and increased disorder of the cytoplasmic domain along with novel transmembrane domain conformations, revealing a mechanistic link between specific lysosomal lipids, transporter regulation, and the enigmatic basis of the ClC-7 slow gate.

## Introduction

Lysosomes function as critical nexuses of metabolic sensing and signaling in addition to their core role in macromolecular digestion. Though the mTOR pathway has been most thoroughly studied in this regard, other signaling pathways emanating from the lysosomal membrane are also fundamentally important to cell homeostasis. Signaling by lysosomal phosphoinositide lipids has recently emerged as an important mechanism in metabolic regulation and disease, with implications for cancer, ALS, and prion disease among others^1–6^. Lysosomal membrane traffic is well-known to be dependent on PI(3,5)P2, with depletion of the lipid causing accumulation of large vacuoles^7–17^. PI(3,5)P2 is also an essential regulator of multiple lysosomal membrane transporters and channels, including TRP-ML1, TPC1/2,, ATP13A2, and ClC-7^18–24^; disruption of PI(3,5)P2 synthesis profoundly impacts lysosomal pH, largely through its action on ClC-7^15,16^. Though structures of some of these lysosomal channels and transporters in complex with PI(3,5)P2 are available, little is known about the molecular mechanisms by which the lipid modulates protein function.

The ClC-7 Cl^-^/H^+^ antiporter is found in lysosomal membranes and in the ruffled border of osteoclasts^25,26^. Though the ion transport function of ClC-7 has been proposed to contribute to lysosomal acidification and to lysosomal Cl^-^ homeostasis, its detailed role remains controversial^25–29^. Loss of function mutations in ClC-7 have long been known to result in osteopetrosis, lysosomal storage, and neurodegeneration but do not alter lysosomal pH^25,30,31^. However, a gain-of-function mutation in ClC-7, Y715C, was recently shown to cause a novel disease, featuring hypopigmentation, organomegaly, and developmental delay; in contrast to loss of function diseases, patients did not have osteopetrosis, but rather a constellation of symptoms including excessively acidic lysosomes, lysosome storage, and accumulated enlarged vacuoles in cells^32^, confirming a role in pH regulation.

Like other members of the CLC family, ClC-7 forms a dimer, with each subunit containing an independent transport pathway residing in its transmembrane domain (TMD), and a cytoplasmic domain (CyD) containing dual CBS motifs^33^. ClC-7 also forms a functional complex with an obligate subunit, Ostm1, which binds to the periphery of each dimer via a single transmembrane helix, with its N terminus forming a luminal cap on the transporter. When driven to express on the plasma membrane^34^, ClC-7 is outwardly rectifying and activates slowly, on the timescale of seconds, at depolarized voltages^35^. Thus, unlike most transporters solely driven by substrate gradients and/or ATP hydrolysis, ClC-7 behaves like a voltage-gated ion channel in that it must be in a voltage-dependent “activated state” for the transport cycle to proceed. CLC family members that act as channels possess a “common gate” that acts on both ion pores simultaneously, in addition to the individual gates in each pore. This is also the case in ClC-7, with the common gate responsible for the slow, voltage dependent activation of the transporter^36^. Although the molecular basis for common gating in the CLC family is unclear, recent studies suggest the interactions between the transmembrane and cytosolic domains play an important part in the gating mechanism.

ClC-7 is inhibited by the lipid PI(3,5)P2, which is generated in the cytosolic leaflet of endolysosomal membranes by the kinase PIKFyve^15^. Depletion of PI(3,5)P2 results in enlarged, hyperacidified lysosomes, with the hyperacidification primarily mediated by ClC-7^15^. The cellular effects of PI(3,5)P2 depletion mimic those found in human patients carrying the Y715C gain-of-function mutation; this mutation prevents the inhibition by PI(3,5)P2, henceforth referred to as “PIP2”, thus apparently allowing increased chloride transport into the lysosome lumen and supporting increased proton buildup.

Recent Cryo-EM structures of wild-type ClC-7 reveal a phosphoinositide bound to the protein with its headgroup facing the cytoplasmic face of the membrane^33^. The negatively charged headgroup of the lipid participates in multiple interactions with the protein; among these is a coordination network that includes the tyrosine residue (715) mutated in gain-of-function patients^32^. Notably, this coordination network includes residues from both the transmembrane domain (TMD) and cytoplasmic C-terminal domains (CyD) of the transporter (the TM-C interface). The interactions between the negative charges on the PIP2 lipid and the complementary positive charges contributed by the protein form an extended electrostatic interface connecting the lipid, TMD and CyD. Here we probe the role of this interface in regulation of ClC-7 by PIP2 using a combination of functional analysis, structure determination, and molecular dynamics simulations.

## Results

### TMD-CyD interactions in slow gating and inhibition by PIP2

To investigate the role of specific interdomain interactions in ClC-7 inhibition by PIP2, we initially examined four residues that contribute to the TMD-CyD-PIP2 interaction network in the WT CLC- 7 cryo-EM structure^33^ (PDB:7jm7): D282, K285, Y715, and R717 (Fig. 1, A-B). We sought to disrupt the charge-charge interactions in this network by introducing D282A, K285E, and R717E mutations individually into a plasma membrane-targeted ClC-7 background characterizing them using whole-cell patch-clamp electrophysiology. Consistent with an important role for the TM-C interface in ClC-7 activation, all three new mutations resulted in gain-of-function phenotypes similar to the effects of the Y715C mutation, including increased current density and faster activation kinetics compared to WT, (Fig. 1, C-D, Fig. S1A-C). Thus, these other interfacial residues are also important to the inhibition mechanism. To assess the integrity of the anion permeation pathway in the Y715C mutant, we replaced extracellular chloride with bromide, iodide, or nitrate. These ion replacements reduced transport currents in a manner qualitatively similar to WT, indicating that current increases in these gain-of-function mutations do not result from substantial changes in the anion pathway (Fig. S1D). All four mutations also abolished or dramatically curtailed the inhibition of ClC-7 by PI(3,5)P2 (Fig. 1C, E), further supporting the idea that the formation of interfacial salt bridges is critical for the inhibition process. Interestingly, a recently identified patient carrying a K285T mutation had a syndrome similar to patients carrying the Y715C mutation, and introducing this mutation diminished sensitivity to PI(3,5)P2 in patch- clamp experiments^37^, confirming our observations and their physiological importance. These two findings, gain-of-function and loss of inhibition, reveal a direct role of the TM-C interface in ClC-7 regulation, and suggest that PI(3,5)P2 inhibits the transporter by inducing formation of this network.

**Figure 1.**
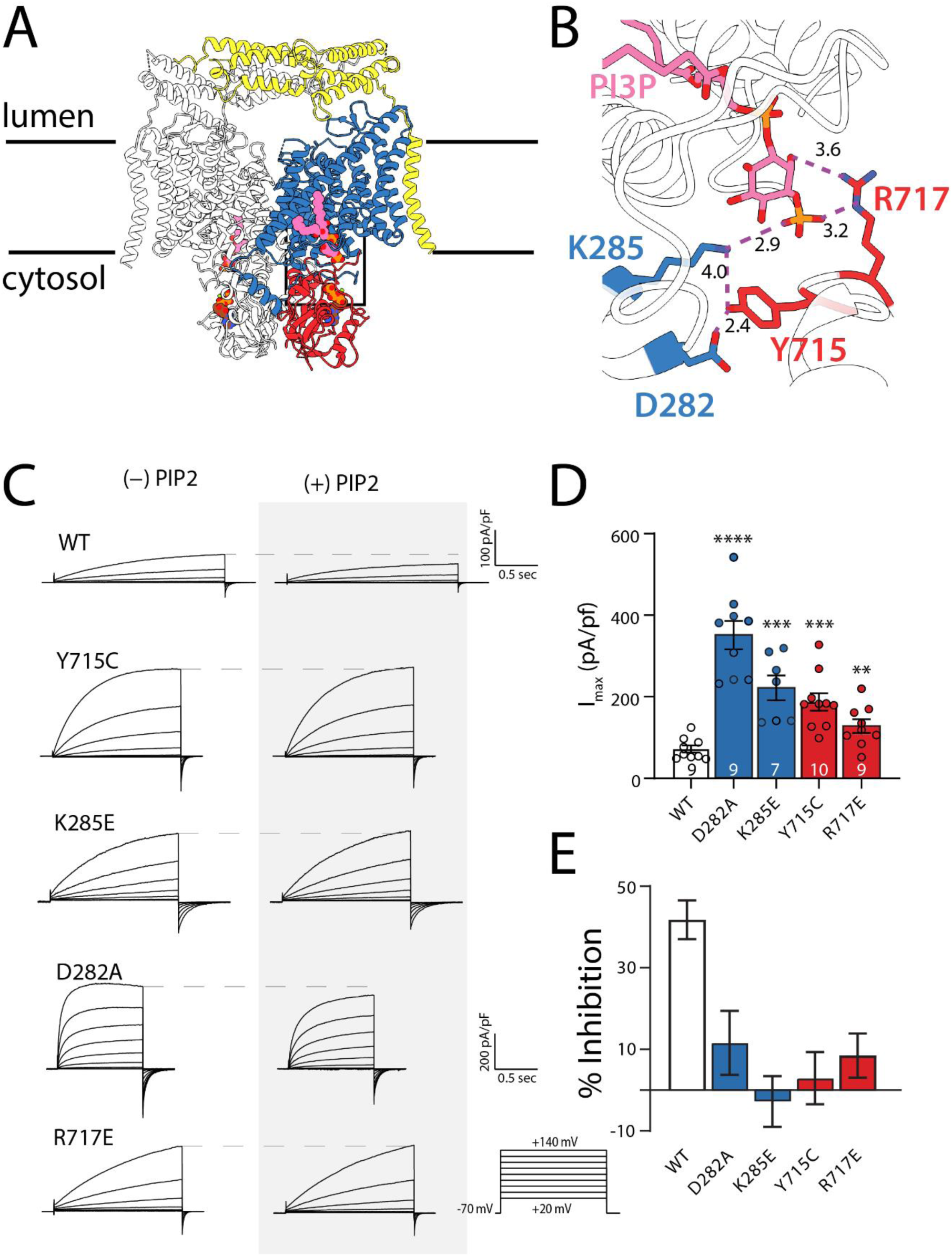
Effects of mutations at the TM/CYD/PIP interface. A) Cryo-EM structure of wild- type ClC-7 (PDB 7JM7) highlighting the bound PI3P molecule. One subunit of the dimer is colored with the transmembrane domain shown in blue, cytosolic domain in red, and Ostm1 in yellow. B) Enlarged view of the interfacial hydrogen bonding network. The phospholipid head group mediates interactions that secure the cytosolic domain to the transmembrane domain. C) Representative traces from whole-cell patch clamp of mutants that disrupt the interaction network in the presence and absence of PI(3,5)P2.. Y-axis scale bar is 100 pA/pF for all traces except D282A. D) Mean Peak currents measured at the end of a depolarizing voltage pulse to +140 mV for TMD (blue) and CyD (red) mutants. Individual experimental points are shown for each mutant. Number of biological replicates is indicated in each bar and statistical significance was calculated from a Student’s *t* test comparison with wild-type ClC-7 (D282A, *p* < 0.0001; K285E, *p* = 0.00013; Y715C, *p* = 0.00018; R717E, *p* = 0.00918). E) Percent inhibition by 50 µM PI(3,5)P2 of maximum currents measured at +140 mV. Values were calculated by subtracting the average percent change in current magnitude in the presence of PI(3,5)P2 from the average percent change in the absence of the lipid and propagating the standard error of the means. Numbers of biological replicates are the same as in (D).

The ClC-7 structure reveals that the loop between helices F and G (the F-G loop) also interacts closely with the glycerol backbone of the modeled PI3P lipid, but does not feature extensive interactions across the TM-C interface (Fig. S2 A)^33^. Several amino acids in this loop form H- bonds with the bound lipid, including S266 and T267. In contrast to the interfacial mutations described above, the F-G loop mutants S266A and T267G retained the slow activation times and current magnitudes of wild type ClC-7 but showed substantially reduced inhibition by PIP2 (Fig. S2, B-E). Taken together, these results suggest that these residues in the F-G loop contribute to binding of PIP2, but not substantially to stabilizing the TM-C interface.

### The ClC-7 CyD is necessary for inhibition by PI(3,5)P2 and for slow gating

Like other members of the CLC family, ClC-7 is a dimer with an independent ion pathway contributed by each monomer^22,33,38–43^. Each pathway functions independently but is also under control of a “common gate” that simultaneously regulates both transport pathways in the homodimer^36^. The common gate of ClC-7 activates slowly, on the timescale of seconds; the origin of this slow activation has been attributed to the cytosolic C-terminal CBS domains^36^. Previous work demonstrated that the ClC-7 CyD need not be covalently linked to the TM domain; either separately injected domain RNAs or transplantation of the ClC-7 CyD to the C-terminus of Ostm1 produced functional transporters^36^. We revisited these results in the context of regulation by PI(3,5)P2 by expressing a truncated ClC-7 lacking the C-domain (Fig. 2A), and therefore eliminating the TM-C interaction network. Patch-clamp experiments reveal that absent its CyD, ClC-7 retains its strong outward rectification, but completely loses its slow activation, instead activating almost immediately upon depolarization, reminiscent of currents observed for the endosome-localized family members ClC-4 and ClC-5 (Fig. 2A-C). Inhibition by PIP2 is also lost in this construct, further supporting a critical role for the TM-C interface (Fig. 2D). Though this result differs from that reported by Ludwig et al.^36^, we note that we measured whole-cell currents from patch-clamped mammalian cells, a quite different system than the *Xenopus* oocytes used in their experiments. Like Ludwig et al, however, upon transplantation of the C-terminal portion of the CyD to the cytoplasmic tail of Ostm1 we observe restoration of the slow activation, though the activation kinetics are still faster than the WT protein. Notably, some PIP2 sensitivity is also restored in this construct (Fig. 2D). The results of our split transporter experiments lend further support to the idea that transport through ClC-7 is suppressed by the formation of the TM-C interaction network. The loss of slow activation accompanying CyD removal hints that the TM-C network is important for the slow gating process itself.^33^

**Figure 2.**
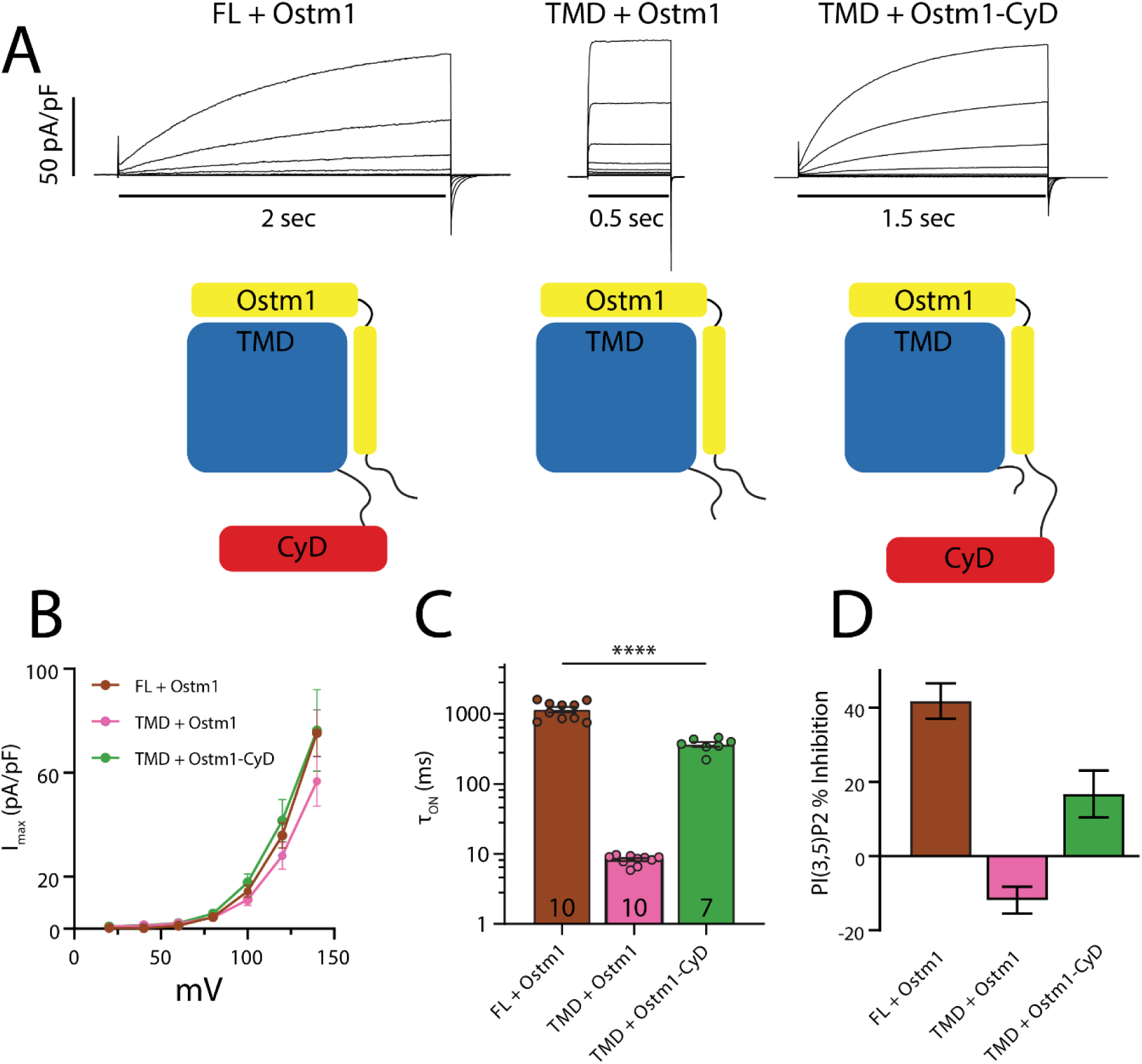
Transporter currents for split ClC-7. A) Representative current families recorded from HEK293 cells transfected with the constructs illustrated below depicting full-length, TMD- only, and split transporter constructs. For simplicity only one protomer is shown. Voltage pulse protocol was the same as shown in Fig. 1C. B) Current-voltage relations from each of the constructs shown in (A) (FL, *n* = 10; TMD+Ostm1, *n* = 10; TMD+Ostm1-CyD, *n* = 7). C) Activation time constants of full-length and split ClC-7 obtained from single exponential fits to currents after +140 mV pulse. Number of biological replicates is indicated in each bar and statistical significance was calculated from a Student’s *t* test comparison with wild-type ClC-7. E) Percent inhibition of maximum currents measured at +140 mV by 50 µM PI(3,5)P2, calculated as described in Fig. 1E. Biological replicates are the same as in (C).

### PI(3,5)P2 induces an ordered CyD

To observe the structural basis of PIP2 inhibition, we used cryo-EM to determine the ClC-7 structure in the presence and absence of PIP2. Since the Y715C mutation completely relieves this inhibition^15,37^, we first determined the structure of this mutant protein. A structure of human ClC-7 Y715C in complex with Ostm1 at 2.6 Å reveals an overall dimeric architecture of Ostm1 and the ClC-7 TM domain quite similar to those of the WT protein (Fig. 3A, Table 1. Fig S3, S4, PDB ID: 9G6E). Remarkably, however, the cytoplasmic domain is completely unresolved in the EM density maps despite its clear presence in our protein preparation, suggesting that the interfacial interaction is not only broken, but that the domain is also either highly mobile or disordered. To control for the possibility that the unresolved domain is due to differences in our preparation from published structures (where the domain is visible and ordered), we solved several further ClC-7/Ostm1 structures following the methods previously described^33^. To our surprise, initial structures of the WT complex also failed to resolve the cytoplasmic domain, despite clear TM domain and Ostm1 density (Fig. 3B, PDB ID: 96GD). However, adding PI(3,5)P2 during purification produced a structure of the WT protein at 1.8 Å resolution that reveals a highly ordered cytoplasmic domain, much as previously reported (RMSD to 7JM7^33^ is 0.858 Å, Fig. 3C, Table 1. Fig. S3, S4, PDB ID: 9G6C); in addition, we also observed density for a bound phosphoinositide. Based on this observation we designate the lipid-free WT structure as the ‘apo’ form of the protein, which we propose corresponds to the active state in the lysosome. Notably, parallel introduction of PIP2 to the ClC-7 Y715C/Ostm1 variant failed to induce the appearance of resolvable density for the cytoplasmic domain. Indeed, of multiple conditions imaged by cryo-EM in the presence of PIP2, several yielded ordered CyDs for WT protein, in contrast to no ordered CyDs in any Y715C structures (Figure S9). Together these results suggest that the pathogenic mutation is sufficient to not only disrupt the TM-C interaction network but lead to complete domain dissociation even in the presence of PI(3,5)P2 (Table 1. Fig S3, S4, S9).

**Figure 3.**
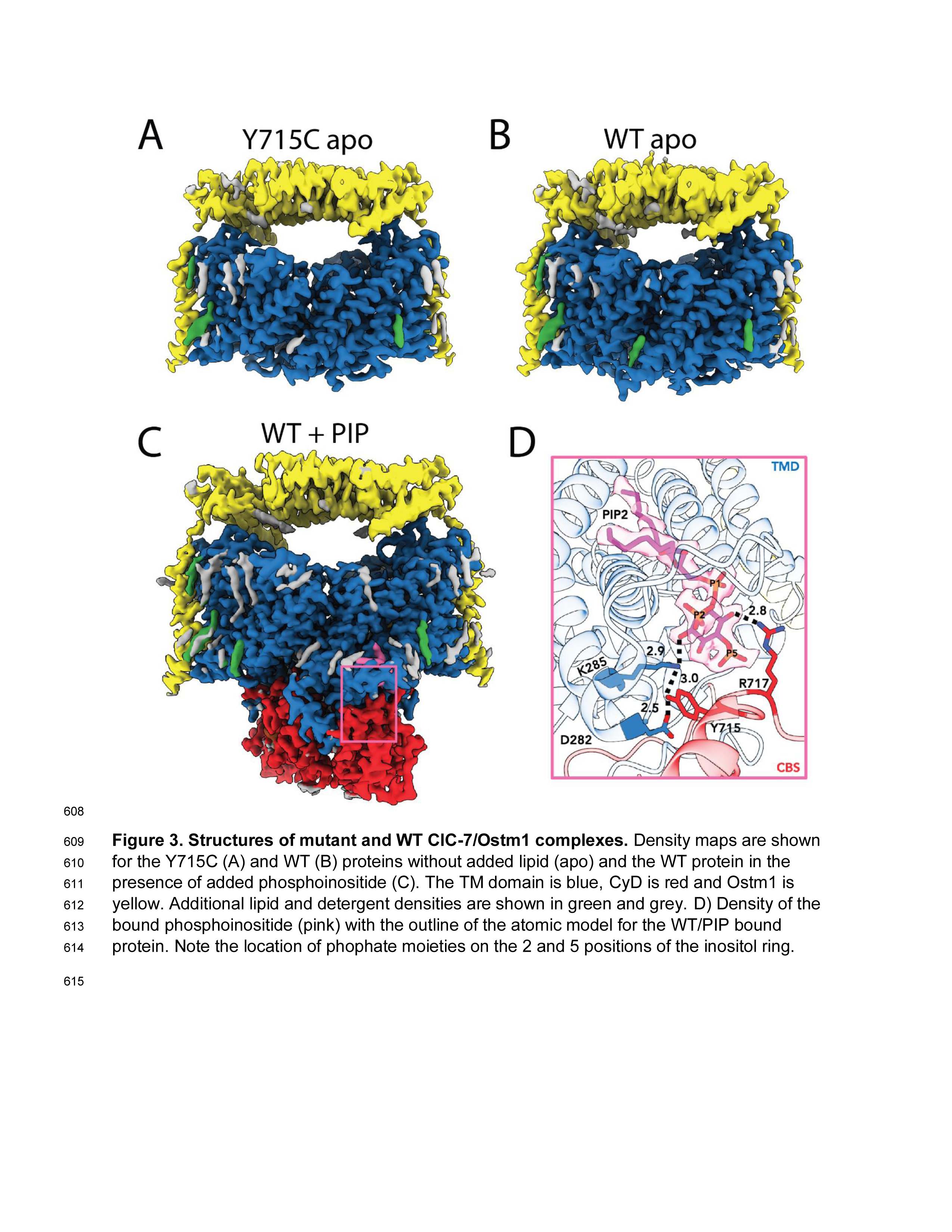
Structures of mutant and WT ClC-7/Ostm1 complexes. Density maps are shown for the Y715C (A) and WT (B) proteins without added lipid (apo) and the WT protein in the presence of added phosphoinositide (C). The TM domain is blue, CyD is red and Ostm1 is yellow. Additional lipid and detergent densities are shown in green and grey. D) Density of the bound phosphoinositide (pink) with the outline of the atomic model for the WT/PIP bound protein. Note the location of phophate moieties on the 2 and 5 positions of the inositol ring.

**Table 1.**
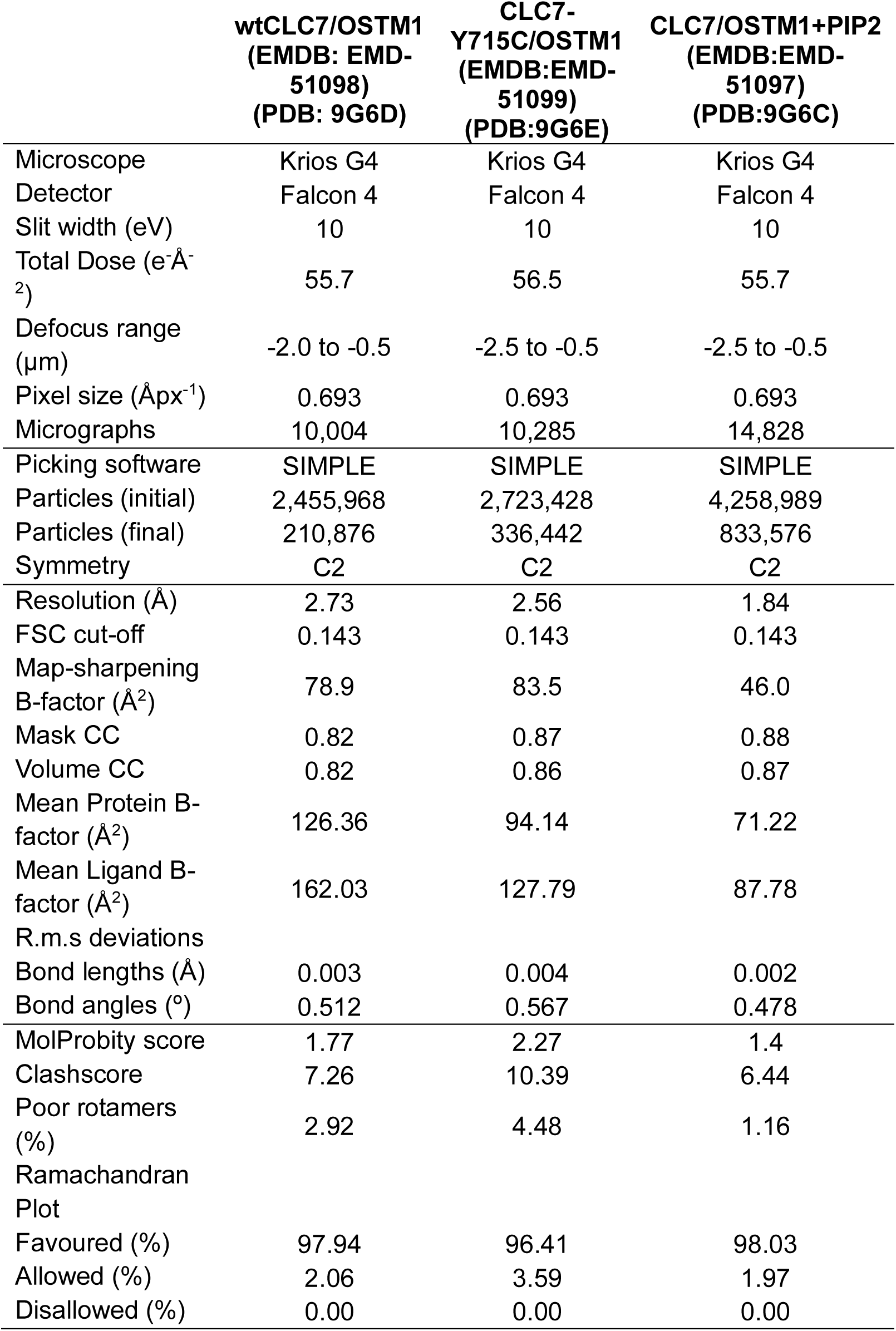
Cryo EM data collection, refinement and validation statistics.

In the WT ClC-7/OstM1 structure the amphipathic loop connecting TM helices F and G extends over the lipid binding site and interacts extensively with the inositol phosphate headgroup and glycerol backbone, sequestering them from the rest of the lipid bilayer (Fig.3 C, D) ^33^. The clear resolution of the F-G loop in lipid-bound structures contrasts sharply with both the apo and Y715C structures, where this loop is poorly resolved, and no clear lipid density appears (Fig S5). The lipid in the CLC-7 structure reported in 2020 was modelled as a PI3P because of stearic clashes if a second phosphate was introduced on the lipid; however, relatively subtle movements of multiple sidechains in our new structure readily accommodate a second phosphate group on the lipid.

Unexpectedly, the high resolution of the new structure clearly reveals the bound lipid to be PI(2,5)P2, a non-physiological phosphoinositide, despite the addition of lipid purchased as PI(3,5)P2 during the purification (Fig. 3D). Chemical analysis reveals that the lipid purchased nominally as PI(3,5)P2 is actually PI(2,5)P2 (Fig S6); however, patch clamp experiments demonstrate that both lipids have similar inhibitory effects on ClC-7 currents (Fig S6), suggesting that insights from this structure also apply to the PI(3,5)P2-bound form. Based on this observation we henceforth refer below to the bound lipid as PIP2.

Taken together, this series of structures and our functional measurements coalesce into a model wherein binding of PIP2 to ClC-7 induces a tight association between TM and C domains, thereby inhibiting transport. In this model, unbinding of lipid or mutation of residues at the interdomain interface weakens the interaction sufficiently to cause domain dissociation even in the presence of PIP2, with resulting increased disorder of the cytoplasmic portion of the protein—an activated state.

### PIP2 induces conformational changes in the ClC-7 TMD

Disruption of the PIP2-TMD-CyD interaction not only causes drastic disordering of the CyD, it also introduces previously unseen conformations of the ClC-7 TMD. While the overall architecture of the TMD in the WT-apo form is varies little from the PIP-bound structure (RMSD 0.539 Å), there is a single major exception: the R-helix (residues 602-609) adopts two conformations (Figure 4), an unprecedented observation in any member of the CLC family. The first conformation resembles that observed in the lipid-bound state and reported previously^33^, with the Q and R helices packing closely to helices D and O and positioning a conserved tyrosine (602 in ClC-7) close to the central chloride permeation pathway, a configuration similar to all previous structures of CLC family members^38^. In this state, which we designate R1, the R-helix forms part of a constriction that limits access of ions to the permeation pathway from the cytoplasmic face of the protein. In the second state, which we designate R2, the R-helix is no longer in that position. Instead, the maps clearly show the unstructured region between Q and R helices (residues 598-600) has itself become helical and formed an extension to the Q helix, which now terminates on the cytoplasmic face of the protein at residue 604, eliminating the R helix altogether. In the new R2 state, the Y602 side chain moves ∼ 15.5 Å away from the chloride binding site, opening a clear conduit from the cytoplasm to the ion permeation pathway (Figure S7). In CLC channels, this tyrosine has been implicated in the common gating mechanism, in which it interacts with the external gating glutamate via motions coordinated through the dimer interface^44,45^. In the bacterial CLC exchanger ClC-ec1, the equivalent tyrosine (Y445) is thought to form an inner gate with a conserved serine (S107) and is critical for the strict 2:1 chloride:proton transport coupling. Mutating this residue in ClC-ec1 leads to decoupled chloride transport, with the degree of decoupling roughly correlated with the size and hydrophobicity of the amino acid^46,47^. Indeed, movements of the R-helix have been suggested to be important in ClC-ec1 function^48,49^. However, the large movements of the R helix seen in our structures have not previously been observed in any CLC structure, and it is unclear whether this movement is part of the transport cycle or part of the unique transport gating of CLC exchangers. Notably, the so-called “proton glutamate^50^” (E314 in ClC-7), which is essential for transport in all CLC antiporters, including ClC-7^51^, is brought into the same aqueous vestibule as the Cl^-^ ions by the conformational change (see discussion).

**Figure 4.**
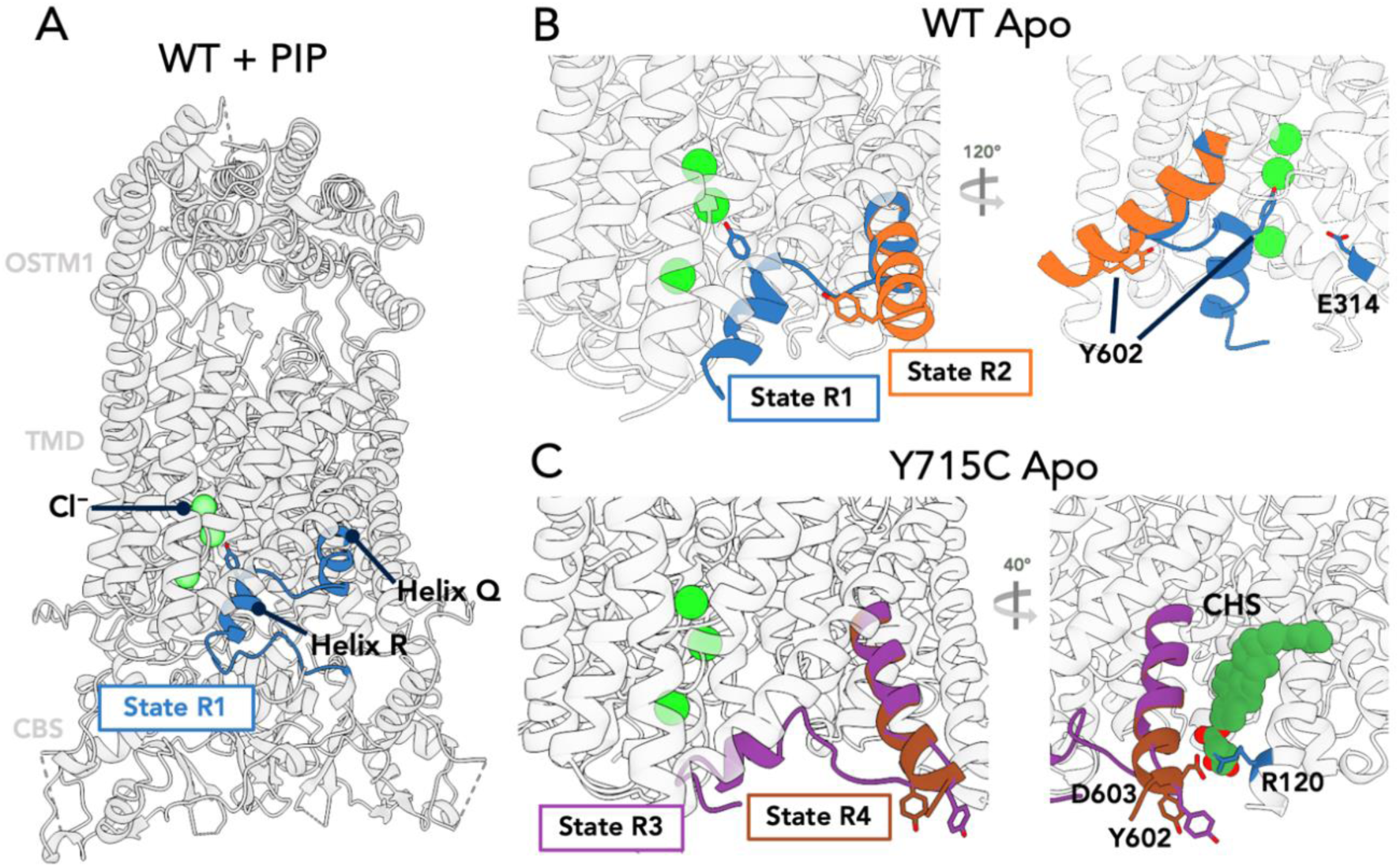
Conformational Heterogeneity in the ClC-7 TM domain. A. WT ClC-7/Ostm1 + PIP2 highlighting helices Q and R (blue) in the R1 state, with Y602 indicated. Green Spheres represent bound Cl^-^ ions. B. WT apo structure showing the two resolved conformations R1 (blue, as in (A)) and R2 (orange), with an extended Q-R helix swinging Y602 to the cytoplasmic face of the protein. C. Y715C apo state showing two additional conformations, R3 (violet) and R4 (brown), both of which show the novel position of residue 602. Not the salt bridge illustrated between D603 and R120 in the right-side view.

Intriguingly, we also observed two additional populations in the cryo-EM maps for the R-helix in the Y715C mutant, which we designate R3 and R4 states (Fig.4 & Fig. S7). In both the R3 and R4 states, the Q-helix has adopted an extended configuration comparable to that observed in the R2 state for the WT protein, with Y602 located at the cytoplasmic interface of the TMD and stabilized through an interaction between D603 and R120 of different protomers, as well as by a bound cholesterol hemisuccinate molecule (Fig.4). The R-helix has essentially unfolded, and a large vestibule is observed opening up from the cytoplasmic side of the membrane to the chloride pathway. The main difference between the R3 and R4 states is that the backbone in the R3 state still contacts helices D and J in the TMD and adopts a short alpha helix, whereas in the R4 state, the Q-helix has extended out and we don’t observe any density for the R-helix region.

Additional changes to the TMD helices are also observed in the Y715C structures. For example, helix J moves approximately 2-3 Å toward the luminal side of the protein, in contrast to the minimal changes observed in the I-J and J-K loop regions (Fig. S7). Helices H, I, and P contribute much of the ClC-7 dimer interface, shifting slightly (RMSD 1.5 Å for H,I and 1.3Å for J) to accommodate the movement of helices R and J. These movements result in the breaking of salt bridges between residues E313 and Q321 on opposing protomers; mutation of these residues was recently reported to affect ClC-7 gating kinetics^52^. Helices L, N, and O also move as a rigid unit toward the center of the protein as a consequence of the conformational shifts of helices J, Q, and R. Overall, the introduction of the Y715C mutation appears to introduce protein rearrangements not seen in lipid-bound or apo WT ClC-7; these changes may underlie the increased activity of the Y715C- mutant transporter.

Despite the numerous changes in the TMD and the dramatic movements of residue Y602, other residues involved in the the Cl^-^ permeation pathway show few substantial differences between our structures (Fig. S7). Residues E247 (the “gating glutamate”), E314 (the “proton glutamate”), S204 and Y505, are all positioned similarly in the WT apo structure, suggesting that the luminal side of the ion conduction pathway is preserved. However, the movement of the R helix in the new conformation substantially changes the cytoplasmic side of the ion pathway for Y715C mutant structure, as shown by a PoreAnalyser analysis^53^ (Figure S7), with substantial changes in the pore-lining residues on this face of the protein.

### Molecular dynamics reveals early steps in interface remodeling

Our structures suggest that in the transport-active apo state of ClC-7, disrupted interaction between the TMD and CyD results in increased conformational dynamics. To test this hypothesis, we performed all-atom molecular dynamics simulations in the presence and absence of the actual PI(2,5)P2 lipid we observed in the structure. We reasoned that simulations starting from the lipid bound state but with the phospohoinositides removed and/or with the Y715C mutation introduced would reveal early steps in the evolution toward the stable states we observe in our structures. To this end, we built a series of systems incorporating ClC-7 and Ostm1 in 100% POPC lipid bilayers starting with our new PIP2 bound structure: we performed simulations with/without PIP2 and with/without the Y715C mutation: for each condition we ran five simulations of 4 µs each, yielding totals of 20 µs per system.

The TMD remained stable over the course of the simulation time for all our constructs (Figure S8) so we focused on the areas of the protein closest to the lipid binding site. As a measure of stability, we calculated the RMSDs of the protein elements engaging the PIP2; specifically the loop region between helix F and G (FG-loop, residues 263-282) which envelops the bound lipid, and the CyD helix containing residue Y715, which we denote α1 (residues 707-715, Figure 5). In the WT PIP- bound trajectories this region was quite stable, with an average RMSD of 1.1 Å; however, the fluctuations of the WT-apo trajectories were substantially larger at 2.0 Å. Similarly, for the pathogenic Y715C mutation, removing the lipid significantly increased the RMSD, to 2.5Å (Figure 5 B, C) with substantial variability even immediately after the equilibration period. We confirmed the flexibility of the apo forms of WT and Y715C by performing parallel simulations using the previously published structure (7JM7) as a starting point (Fig. 5 C, Fig. S8). Consistent with our model and even in the relatively short microsecond timescales of our MD simulations, we observe trends toward increased molecular disorder in both apo and Y715C proteins, but not in the PI(3,5)P2-bound WT model. We predict that longer simulations will reveal further progress toward full domain dissociation.

**Figure 5.**
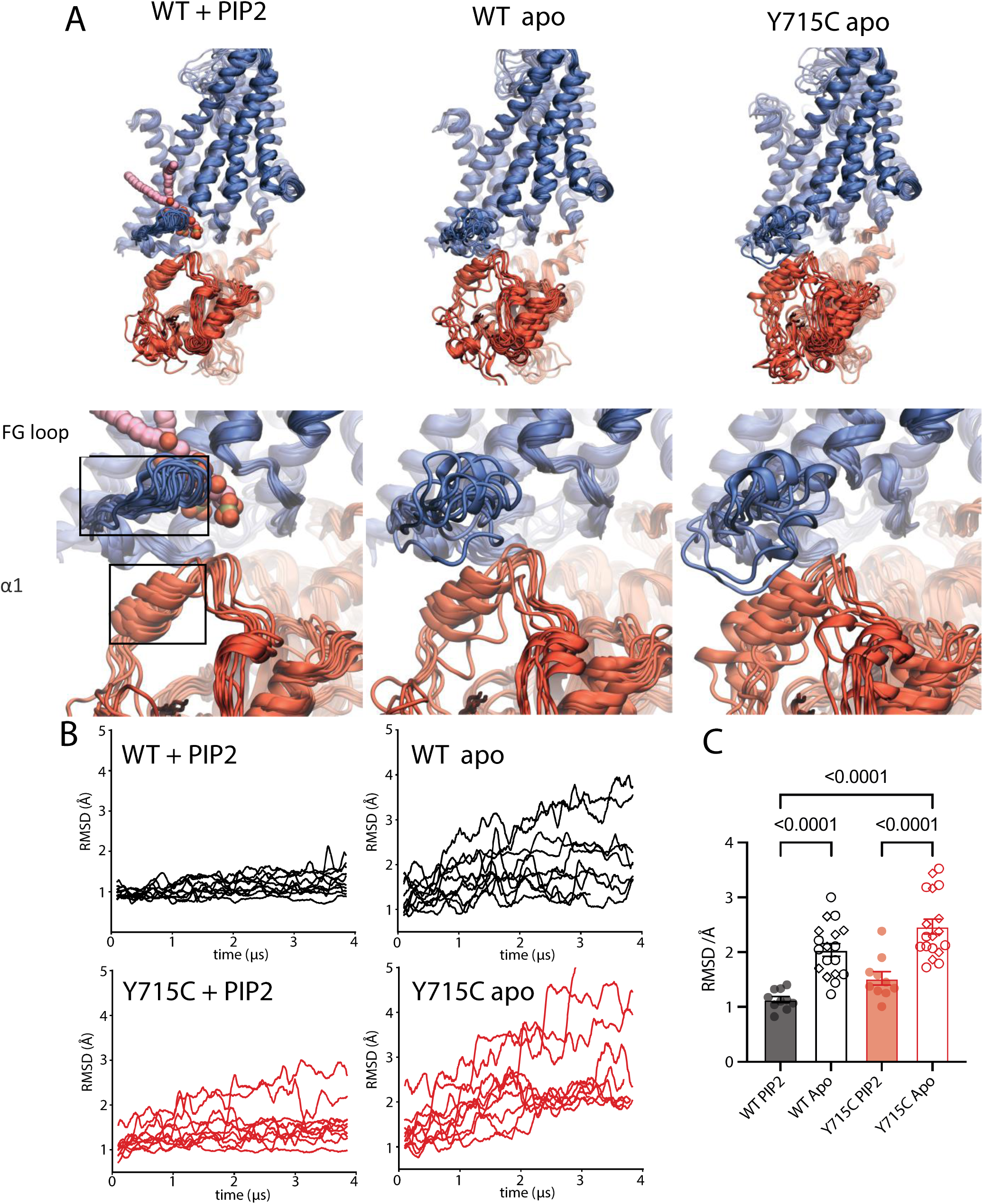
Molecular dynamics simulations reveal effect of PIP2 on F-G loop and. α1 **motions.** A. Snapshots illustrating the flexibility of the FG loop (residues 263 – 282) and ⍺1 (resid 707 – 715) region in various simulation states: WT in the presence of PIP2 (left), WT apo (center), and Y715C apo (right). Each protomer from the dimer in all five replicates is aligned onto a single protomer from the 7JM7 cryo-EM structure. Bottom row is a zoom-in of the snapshots from the top row (WT PIP2-bound, left; WT apo, middle; and Y715C apo, right). Boxed in the bottom row (left) are the FG loop and helical region α1. B. Average Root Mean Square Deviation (RMSD) time series for FG loop and ⍺1 regions from WT/WT apo (black traces) and Y715C/Y715C apo (red traces). RMSDs were calculated relative to the FG loop (residues 263 – 282) and α1 (residues 707 to 715) regions in the cryo-EM structure, and each trace was calculated from one protomer. C. Per protomer average RMSDs of FG loop and ⍺1 regions calculated as in B, but excluding the initial 500 ns of the trajectory as equilibration. Empty circles or diamonds are used to represent apo protomers from our structure or PDB 7JM7 respectively. Adjusted p-values were calculated using an ordinary one-way ANOVA and Tukey test for multiple comparisons. Statistically significantly different columns are indicated; all other comparisons were not significant.

## Discussion

Our results lay out a clear model of PIP2 inhibition of ClC-7 and explain the gain-of-function resulting from pathogenic mutations identified in human patients. Briefly, binding of PI(3,5)P2 induces a strong interaction network between charged residues in the TMD and CyD of ClC-7, stabilizing order in the CyD and forcing association with the TMD. This highly stable state precludes the conformational changes necessary for transport. Either unbinding of PIP2 in WT ClC-7 or introducing mutations that disrupt the network destabilizes the TMD-CyD association and allows for the conformational flexibility in the Q-R helices necessary for Cl-/H+ exchange (Fig. 6).

**Figure 6.**
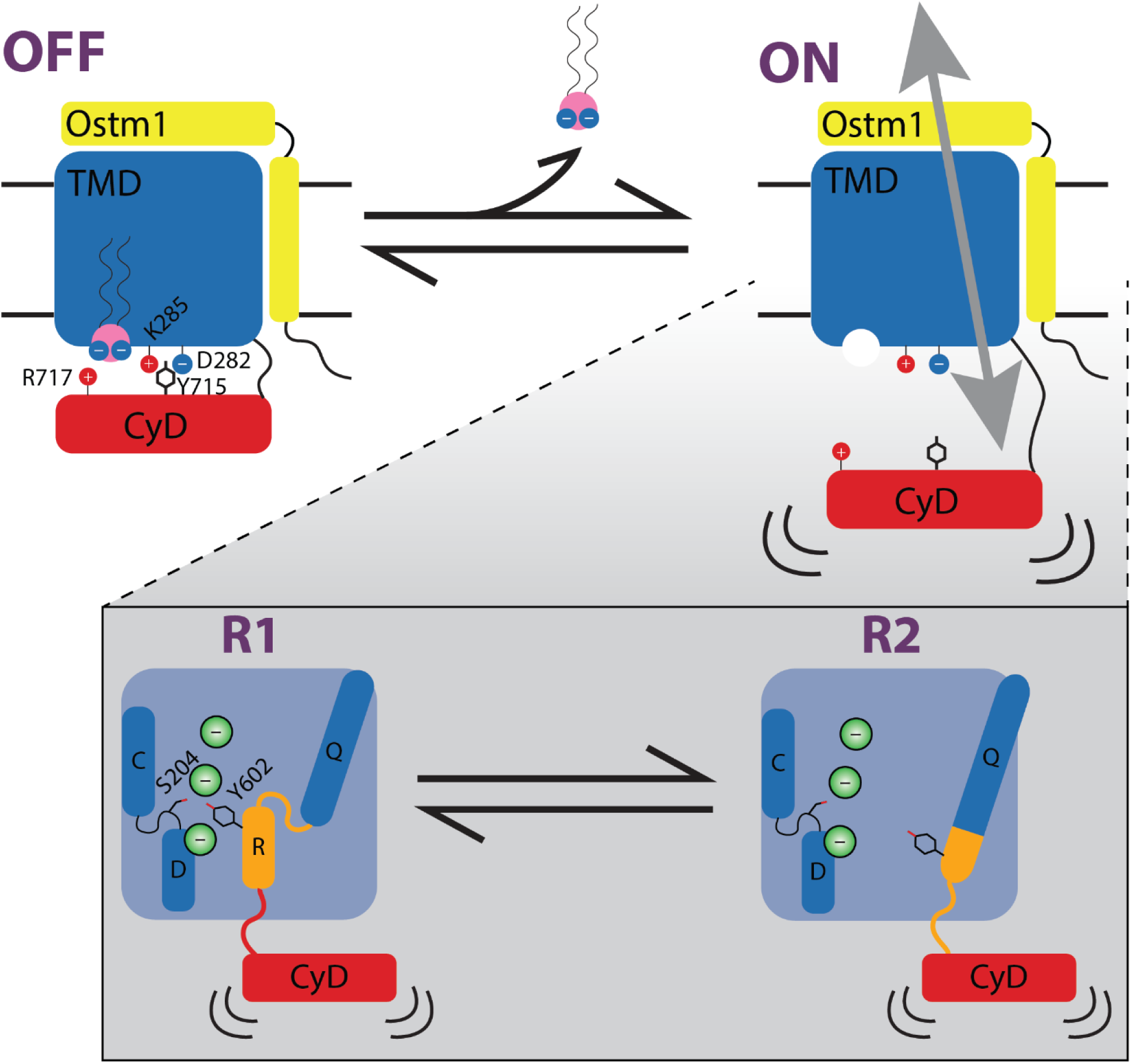
Proposed model of ClC-7 regulation by PI(3,5)P2. In the PIP-bound state (OFF), lipid binding creates an optimal interface to stabilize and anchor the ordered CyD against the TMD, with key residues highlighted. In the absence of PI(3,5)P2 (ON), this interface is lost and the CyD becomes disengaged and dynamic. Within this apo state, loss of the stabilizing interactions allows for the substantial movement of the critical R helix observed in the PIP-free or Y715C mutant structures (R1 and R2). Whereas the Q-R helix in the R1 conformation is similar to the PIP-bound, inactive structure, in the R2 conformation the ion transport pathway becomes much more solvent-accessible. These two conformations may represent steps along the gating or the transport pathway.

We present three main lines of evidence to support this model. First, mutating residues within interaction distance of the charged headgroup of the phospholipid has dramatic functional consequences in electrophysiology experiments: faster activation, larger currents, and loss of PI(3,5)P2 inhibition. Mutating residues in the F-G loop, which contribute to PIP binding but not to the domain interface, retains wild-type-like currents but dampens the effect of PIP2, consistent with a weakened binding site but retained domain interface. Completely removing the cytosolic domain results in very fast activation with accompanying loss of inhibition, both partially restored by fusing the cytosolic domain to the nearby cytosolic C terminus of Ostm1.

Second, our high-resolution structures reinforce the idea that PIP2 is necessary to stabilize the TMD-CyD interface via a network of electrostatic interactions. When PIP2 is not present in the sample, the CyD is unresolvable, suggesting that it becomes unbound from the TMD and highly dynamic. Further, when the pathogenic gain-of-function mutation Y715C is introduced, PIP2 is unable to to induce rigid binding of the C-domain to the TM domain. This comports with our previous results showing that PIP2 has no measurable effect on the mutant in electrophysiology measurements^15^. Our data currently do not distinguish whether the unresolved CyD in our structures results from a high mobility of the CyD compared with the TMD or from a complete unfolding of the domain. Further experiments will be required to distinguish these possibilities.

Finally, all-atom molecular dynamics simulations of full-length ClC-7 show that in the presence of the Y715C mutation or in the absence of PIP2, parts of the CyD rapidly destabilize, even on the microsecond timescale of our simulations, setting the stage for the larger conformational changes which occur over the seconds timescale too long to simulate with current methods.

Though our focus here is the mechanism of PIP inhibition of ClC-7, the acute effects of interface mutations on the kinetics of the slow, voltage-dependent activation process and the profound gating acceleration observed in the TM-only transporter hint that slow gating and PIP2 inhibition are at least tightly coupled, if not part of the same process. In addition to loss of inhibition, mutation of most residues in the TMD-PIP2-CyD network leads to faster activation and slower deactivation of ClC-7 currents. This is expected in the context of our simplified model, in which the activation time constant reflects unbinding/disordering (off-rate) of the CyD to the TMD and deactivation time constant reflects re-binding of the CyD (on-rate). In the gain-of-function mutants, the ratio of these rates is shifted to favor the activated state of the transporter (Fig. S1 C). This contrasts with the effects of mutating residues in the F-G loop, which minimally affect gating kinetics. Thus, rather than completely blocking activation of the transporter, PIP2 binding may shift the rates to favor the inactive state, a possibility consistent with the partial inhibition we attained in electrophysiology experiments. The dramatic increase in the rate of activation in our TMD-only construct reinforces the importance of the C-domain in the slow voltage-dependent gating of ClC-7. This conclusion clarifies and provides a mechanistic underpinning for the extensive work that has implicated the cytoplasmic domain in the slow gating process in other CLC channels and transporters^36,42,54–56^. We suggest that parallel mechanisms of CyD binding/unbinding to the TM could underlie slow gating in these other proteins. Our conclusions are also fundamentally consistent with those recently reported for the bacterial CLC, ClC-ec1^49^. In that work major rearrangements of cytoplasmic helices facilitate transitions to the active form of the protein and seem to foster further, more subtle rearrangements of helices at the dimer interface as we observe in the Y715C structure. However, the relationships between pH- and Cl^-^ -dependent gating in the bacterial transporter and voltage-dependent gating in ClC-7 remain to be determined.

The novel conformations we observe in the ClC-7 TMD raise intriguing possibilities about the activation and transport cycle of ClC-7. We note that the traditional R-helix is present in the PIP- bound, inhibited WT protein and in the R1 conformation of the apo WT ClC-7, but is absent from both R3 and R4 conformations of the Y715C mutant, which represents the perhaps maximally activated transporter. The extended Q helix in the Y715C mutant has moved even farther from the chloride transport pathway compared to WT apo. In the R4 conformation of Y715C, the Q-R loop is longer, resulting in a shorter R helix in the chloride pathway and comprising residues different from the R helix in WT ClC-7. These observations suggest at least two possible interpretations. It may be that we are observing steps along the activation landscape, with the canonical R helix representing an ‘inactive’ state of the transporter and the alternate extended Q conformation as the active state; in this case the other more subtle differences between the mutant and apo WT structures are less important for gating. Alternatively, the two conformations of the R helix could represent different states in the transport cycle. Further work will be required to distinguish these possibilities.

A long term challenge in understanding CLC transport mechanisms has been the extended hydrophobic region between sites required for proton transport^46^. Experiments and simulations have suggested the possibility of a contribution from the conserved tyrosine (Y445 in ClC-ec1, Y602 in ClC-7)^46^ or an extended water wire^57,58^. Our new structures raise new possibilities. The conformational change to the R2 state in our WT apo structure opens up a previously unseen cavity on the cytoplasmic side of the protein that brings the inner proton transport site^50^ into an aqueous vestibule that extends to the Cl- conduction pathway, eliminating the large separation between proton binding sites (Fig S7). We speculate that the R2 state represents a previously unseen “active” state of the transporter.

Given that ClC-6 and ClC-7 share ∼44% sequence identity, including most of the residues we identify here as critical to inhibition, and similar lysosomal localization, it seems likely that PI(3,5)P2 would have a similar effect on ClC-6. A recently published structure of ClC-6 reveals a protein in a highly similar conformation to the PIP-bound ClC-7^59^. Indeed, inspection of the ClC- 6 maps reveals density consistent with a bound PIP in a similar location and pose to the one we observe.

In contrast, although PI(3,5)P2 also inhibits the endosomal CLCs, ClC-3, 4 and 5, the mechanism of this action appears to be strikingly different^60–62^. In the case of the endosomal exchangers, a separate subunit, the single-span membrane protein TMEM9, directly binds to ClC-3. This results in a tripartite interaction between phosphates in the lipid, residues in transmembrane helices in ClC-3, and residues in the cytosolic domain of TMEM9^62^. When both PI(3,5)P2 and TMEM9 are bound to ClC-3, the cytosolic helix of TMEM9 occludes entry to the chloride transport pathway without noticeable changes in the ordering of the CyD, a markedly different mechanism of inhibition compared with ClC-7. Although PI(3,5)P2 binds to the face of ClC-3 homologous to the PIP binding location we report here for ClC-7, the cytosolic domains of ClC-3 do not appear to interact with its transmembrane domain in proximity to the bound PI(3,5)P2. Moreover, TMEM9 appears to contain phosphorylation sites that may be involved in regulation of inhibition^61,62^. Thus, the endosomal exchangers are likely subject to more layers of regulation to effect inhibition.

PIP regulation of ClCs also appears to span both animal and plant orthologs: a structure of the Arabidopsis atClC-a, also regulated by phosophoinositides, shows a different PIP, PI(4,5)P2, bound at a different location adjacent to the dimer interface^63^. Even in that structure, though, extensive electrostatic interactions between the PIP headgroup and charged residues on both the TMD and CyD raise the possibility that engagement of the domains plays a similar regulatory role in that system even in the context of a completely different lipid binding site.

These findings together with our data establish a general paradigm in which ClC activity must be tightly controlled to prevent excessive acidification and/or chloride accumulation in endolysosomal compartments. Nevertheless, the signals, effectors, and mechanisms that regulate cellular levels of PI(3,5)P2 inhibition remain largely unknown. An emerging player may be SNX10, a protein involved in membrane trafficking that appears to regulate ClC-7 activity by influencing phosphoinositide levels, though the details of this process are still unclear^64^. Our work adds another node to the dynamic web of lysosome regulation and the multiple disparate roles of PI(3,5)P2. PI(3,5)P2 directly regulates pH via the V-ATPase itself by binding to the a-subunit and promoting V_1_-V_o_ assembly^65,66^. Thus, PI(3,5)P2 may have opposite effects on the V- ATPase and ClC-7, with the former promoting acidification and the latter putting the brakes on it. The balance between these and other PI(3,5)P2-dependent processes likely determines the pH attained at a given abundance of the lipid. Beyond pH, endolysosome trafficking and fusion are also regulated by the lipid via PI(3,5)P2-dependent TRPML1 calcium release^67^. TPC1 and 2 channels are also activated by PIP2, resulting in calcium and/or sodium release which impinges on endolysosomal trafficking and autophagy^68^. Finally, PI(3,5)P2 is an upstream regulator of the mTORC1 complex, in which the Raptor subunit binds directly to the lipid to regulate the localization of the complex^69,70^. Thus, PI(3,5)P2 is a nexus of lysosome regulation at multiple levels: pH, trafficking, and metabolic control of the cell.

## Methods

### Electrophysiology

HEK-293 (ATCC CRL-1573) cells were cultured in 35 mm tissue-culture treated polystyrene dishes (Falcon) at 37°C in the presence of 5% CO_2_. The growth medium was Dulbecco’s modified Eagle’s medium supplemented with 10% FBS, 100 units mL−1 of penicillin-streptomycin, 4 mM L-glutamine, and 1 mM sodium pyruvate (Gibco). Cells were transiently co-transfected with 800 ng of human wildtype or mutant ClC-7 in a pIRES2–EGFP vector and 700 ng human Ostm1 in a pCMV-6 vector (for a final 1:1 molar ratio of plasmids). All ClC-7 constructs used for electrophysiology also included mutations to two lysosome-targeting motifs in the N terminus, LL23/24AA and LL68/69AA, to cause trafficking to the plasma membrane^34,36^, allowing for whole-cell measurements. All mutations were introduced using a Q5 Site-Directed Mutagenesis kit (New England Biolabs) and verified by Sanger sequencing. Transfection was accomplished using Lipofectamine 3000 with P3000 reagent (Invitrogen) and performed approximately 40 hr prior to experiments.

To prepare cells on the day of patch-clamp experiments, cells were lifted from culture dishes by brief exposure to 0.25% trypsin/EDTA, resuspended in supplemented DMEM, plated on glass coverslips, and incubated for 1–2 hr at 37°C in 5% CO_2_. Patch pipettes were pulled with a P-97 puller (Sutter Instruments) from borosilicate glass capillaries (World Precision Instruments) and heat-polished using an MF-200 microforge (World Precision Instruments). Pipette resistance was 2–5 MΩ in the extracellular solution. A reference electrode was placed in a separate chamber containing 1 M KCl and connected to the bath solution via a 2% agar bridge made from extracellular solution. Whole-cell voltage-clamp current measurements were performed using an Axopatch 200B amplifier (Axon Instruments) and pClamp 11.1 software (Axon Instruments). Data were acquired at 10 kHz and filtered at 1 kHz. Extracellular solution consisted of (in mM) 130 NaCl, 5 KCl, 1 MgCl_2_, 1 CaCl_2_, 20 HEPES, and 10 glucose, with the pH adjusted to 7.4 using NaOH and the osmolality adjusted to 310 mOsmol with sucrose. For chloride substitution experiments, NaCl was replaced with the same concentration of the sodium salt of the given ion. Pipette solution contained (in mM) 110 CsCl, 10 NaCl, 3 MgCl_2_, 0.5 CaCl_2_, 1 EGTA, and 40 HEPES; pH was adjusted to 7.2 with CsOH, and osmolality was adjusted to 300 mOsmol using sucrose. Chemicals were purchased from Sigma. Osmolality was measured using a Vapro 5600 vapor pressure osmometer (Wescor). For measurements including PI(3,5)P2, short-chain PI(3,5)P2 diC8 (Echelon Biosciences #P-3508 or Avanti Polar Lipids #850184) was dissolved in water at a concentration of 2 mM and stored at –80°C in small aliquots; on the day of experiments, this stock PIP2 was diluted into pipette solution for a final concentration of 50 µM and kept on ice throughout the experiment.

### Construct Cloning for Structural Biology

Gene encoding Homo Sapiens CLC7 was synthesised, and subcloned into a pLexM mammalian expression vector with a C-terminal 6His-tag, Y715C mutation was introduced with quickchange mutagenesis. Gene encoding Homo Sapiens OSTM1 was synthesised, and subcloned into a pLexM expression vector with a C-terminal Flag-tag.

### Protein Expression in HEK293F Cells and Purification

For expressions in HEK293F cells, equal amounts of plasmids of CLC7-His (or CLC7Y715C- His) and OSTM1-Flag were mixed with PEI Max at a 1:2 ratio for 15 minutes and then used to transfect Freestyle 293F cells (Thermo Fisher Scientific). Sodium Butyrate was also added at the time of transfection to a final concentration of 10 mM. 40 hours after transfection, cells were harvested by centrifugation and stored at -80 °C. Thawed cells were then resuspended in phosphate-buffered saline (PBS) with DNaseI (Sigma-Aldrich), homogenised by sonication, and cell debris was removed by centrifugation. Membrane fractions containing expressed protein were then pelleted by ultracentrifugation, washed with 15 mM Hepes buffer at pH7.5, with 20 mM KCl, and palleted again, before being resuspended in PBS and stored at -80 °C.

For the purification of CLC7/OSTM1 complex, membrane was thawed and solubilised in 1% (w/v) LMNG, 0.1% CHS, 10% Glycerol, and 150 mM NaCl. Solubilised proteins were separated from other fractions by ultracentrifugation, followed by binding by gentle stirring to Pierce Anti-DYKDDDDK affinity resin (Thermo Fisher Scientific) for 2 hours in buffer equilibrated with PBS, 1% LMNG, 0.1% CHS, 10% glycerol, 150 mM NaCl, (Buffer A). FLAG affinity chromatography was then performed in an Econo-Column (Bio-Rad) by washing with 10 column volumes of Buffer A, 30 column volume of Buffer A with 10 mM ATP, 10 mM MgCl_2_, another 10 column volume of Buffer A, with addition of 50 mM PI(3,5)P2 (Avanti) in samples with PIP2, and finally eluting in 5 column volume of Buffer A with 0.5 mg/ml flag peptide (Merck millipore). Flow through collected from the first flag purification was mixed with additional FLAG resin and subjected to another 1.5 hour of rotation, after which transferred to gravity column, washes and elutions were repeated. Elutions containing CLC7/OSTM1 complex were concentrated (Amicon, Millipore, 100 kDa molecular weight cut off (MWCO)) to 500 ul and subjected to size exclusion chromatography in Superdex 200 Increase 10/300 GL column (GE Healthcare). Main fractions containing the complex were pooled and concentrated.

### Structural Studies by Cryo-Electron Microscopy

CLC7/OSTM1 complexes were concentrated to 0.75-1 mg/ml and the CLC7/OSTM1 + PIP2 sample was supplemented with 0.1 % LMNG and 2.5 mM ATP prior to vitrification. Complexes were adsorbed to glow-discharged holey carbon-coated grids (Quantifoil 300 mesh, Au R1.2/1.3) for 10 s, blotted for 2 s at 100% humidity at 10 °C, and then plunge frozen in liquid ethane using a Vitrobot Mark IV (Thermo Fisher Scientific).

Data were collected in counted mode in Electron Event Representation (EER) format on a CFEG-equipped Titan Krios G4 (Thermo Fisher Scientific) operating at 300 kV with a Selectris X imaging filter (Thermo Fisher Scientific) set to a slit width of 10 e-V at 165,000x magnification on a Falcon 4 direct detection camera (Thermo Fisher Scientific), with a physical pixel size of 0.693 Å. Movies were collected at a total dose of 55.7 – 56.5 e-/Å2 fractionated to ∼ 1 e-/Å2 per frame.

Patched (20 x 20) motion correction, CTF parameter estimation, particle picking, extraction, and initial 2D classification were performed in SIMPLE 3.0 ^71^. All downstream processing was carried out in cryoSPARC 3.3.1 ^72^, except for Bayesian particle polishing which was performed in RELION 3.1 ^73^ using the csparc2star.py script within UCSF pyem ^74^ to convert between formats. Global resolution was estimated from gold-standard Fourier shell correlations (FSCs) using the 0.143 criterion and local resolution estimation was calculated within cryoSPARC. For WT-CLC7/OSTM1 and CLC7Y715C/OSTM1 (supplementary Figure 3), multiple rounds of 2D classification were performed followed by multi-class ab initio reconstructions (k=4 or k=5) in C1. Particles belonging to the strongest class were selected and subjected to non-uniform refinement against their corresponding volume lowpass-filtered to 30 Å with C2 symmetry to yield a 2.9 Å volume. Aligned particles were imported into RELION for Bayesian polishing. Polished particles were subjected to non-uniform refinement in cryoSPARC with per-particle defocus, tilt, trefoil, spherical aberration, and tetrafoil estimation/correction applied to generate the final volumes (2.6-2.7 Å).

For WT-CLC7/OSTM1 with PIP2 (supplementary Figure 3), multiple rounds of 2D classification were performed followed by multi-class ab initio (k=4) in C1 with a subset of particles. The resulting volumes, lowpass-filtered to 20 Å, were then used as references for heterogenous refinement (k=4) in C1 with the full particle set. Particles belonging to the strongest volume and most dominant class were selected and subjected to non-uniform refinement in cryoSPARC with C2 symmetry to yield a 2.8 Å volume. Aligned particles were then imported into RELION for Bayesian polishing. Following an additional round of 2D classification, selected particles were subjected to non-uniform refinement in cryoSPARC, yielding a 2.0 Å volume. An additional round of heterogeneous refinement against volumes generated from multi-class ab initio reconstructions (k=4) of a subset of polished particles was performed to further purify the dataset. Selected particles were further non-uniform refined after per-particle defocus estimation to yield a 2.0 Å volume which could be further improved to 1.8 Å after tilt, trefoil, spherical aberration, and tetrafoil fitting.

### Model Building and Refinement

The structure of hsCLC7/OSTM1 (PDB:7JM7)^33^ was manually docked into the wtCLC7/OSTM1 PIP2-bound density map using UCSF ChimeraX^75^. The model was then built according to the density using Coot^76^. Atomic coordinates were refined iteratively against un-sharpened map with geometric and Ramachandran restraints with PHENIX real space refinement^77^.

The refined structure of wtCLC7/OSTM1-PIP2 was then docked into the wtCLC7/OSTM1 apo and CLC7-Y715C/OSTM1 density maps respectively with UCSF ChimeraX, and built according to the density, with geometric and Ramachandran restraints, using Coot, followed by iterative PHENIX real space refinement against un-sharpened map.

### MD simulations

Initial coordinates for human ClC-7 in complex with OSTM1 and ATP, including bound magnesium and chloride ions were taken from the protein data bank (ID 7JM7^33^) and our new PI(2,5)P2-bound WT structure. Missing residues were modeled using the SWISS-MODEL server^78^. PROPKA 2.0^79^ was used to predict the protonation state of ionizable groups. Notably, the so-called gating glutamate (E247) was protonated. Proteins were subsequently embedded in a pure POPC membrane and surrounded with TIP3P water using PACKMOL-Memgen^80^. For apo systems, we removed PI(3)P or PI(2,5)P2 from 7JM7 and our WT structure respectively before running it through PACKMOL-Memgen to ensure no vacuum had been left in our system. We used the mutagenesis tool in PyMOL (The AxPyMOL Molecular Graphics Plugin for PowerPoint, Version 3.0 Schrödinger, LLC.) to generate Y715C mutant from the wild-type structures. Six different constructs were simulated altogether; WT apo from either 7JM7 or our structure, Y715C from either 7JM7 or our structure and PI(2,5)P2 bound WT or Y715C mutant. Each construct was simulated in five replicates for 4 µs, yielding 120 µs aggregate simulation time.

Our simulation box had dimensions of 147Å x 147Å x 163Å with a total of 568 POPC lipids, 81100 TIP3P water molecules and 150 mM KCl. Amber ff14SB^81^ and Amber lipid17^82^ force fields were used for protein and lipid respectively. Parameters for PI(2,5)P2 were obtained from the General Amber Force Field (GAFF2)^83^, while those for ATP were obtained from the Amber parameter database of the Bryce Group at the University of Manchester (http://amber.manchester.ac.uk). Each system was equilibrated with a 25kcal/mol/Å^2^ restraint on heavy atoms which was eased over 120ps and followed by a 20ns unrestrained equilibration. Berendsen^84^ and Monte Carlo barostats were used during equilibration and production respectively, while temperature was kept at 300K with a Langevin thermostat, and a friction coefficient of 1 ps^-1^. Long-range electrostatic interactions were treated using the Particle Mesh Ewald method^85^. Leonard-Jones forces were switched smoothly to zero in the range 10-12Å. We used semi-isotropic pressure coupling for all equilibration and production simulations. The SETTLE algorithm was used to keep water molecules rigid while the SHAKE algorithm was used to convert bonds to hydrogen into rigid constraints. Hydrogen masses of heavy atoms were repartitioned to 3.024 Daltons and production simulations run with a 4fs time step in Amber22 (D.A. Case, Amber 2022, University of California, San Francisco). We generate molecular graphics using the Visual Molecular Dynamics (VMD)^86^ and PyMOL software packages.

RMSD analyses were done using LOOS^87^ and in-house scripts. We used alpha carbons of helical regions in TMD RMSD calculation, while alpha carbons of residues 263 – 282 (FG loop) and 707 – 715 (α1) were used to measure flexibility of the TMD-CyD interface. The corresponding residues from WT PI(2,5)P2 were used as reference in all RMSD calculations.

## Acknowledgements

This work was supported by the NINDS intramural program (JKH, ES, JAM), MG funding source: NIH R01 GM137109 (MG), Wellcome Trust, (YL, JP, SN), and the NCI Intramural Program (JD, SL), MJL is a Royal Society University Research Fellow. The contributions of the NIH author(s) are considered Works of the United States Government. The findings and conclusions presented in this paper are those of the author(s) and do not necessarily reflect the views of the NIH or the U.S. Department of Health and Human Services.

