## Supplemental Figures for "Mechanism of phosphoinositide regulation of lysosomal pH via inhibition of CLC-7"

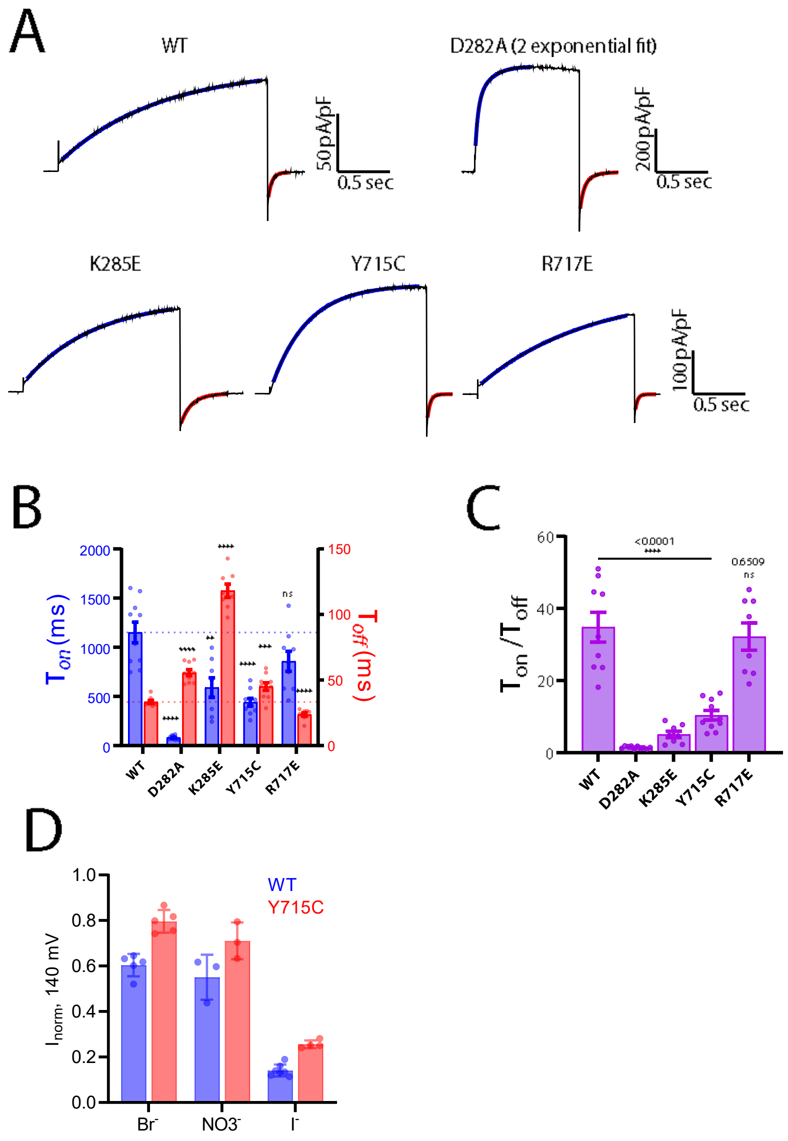


**Supplementary Figure 1. Effect of mutations to the PIP2 binding site on transporter gating kinetics.** A) Representative traces and exponential fits to activation and deactivation of transport currents. Currents from the D282A mutant were better fit by a two exponential function; the slow component values are used in later panels. B) Activation and deactivation time constants extracted from exponential fits, with statistical significance calculated from a Student's *t* test comparision with wild-type (WT, $n = 10$; D282A, $n = 10$, $p < 0.0001$; K285E, $n = 8$, $p = 0.0016$; Y715C, $n = 10$, $p < 0.0001$; R717E, $n = 9$, $p = 0.0617$) . C) Ratio of activation to deactivation time constants; a smaller ratio increased active-state stability. D) Maximum currents measured after replacing external chloride with the indicated anion. Currents were normalized to those measured in chloride solution from the same cell.

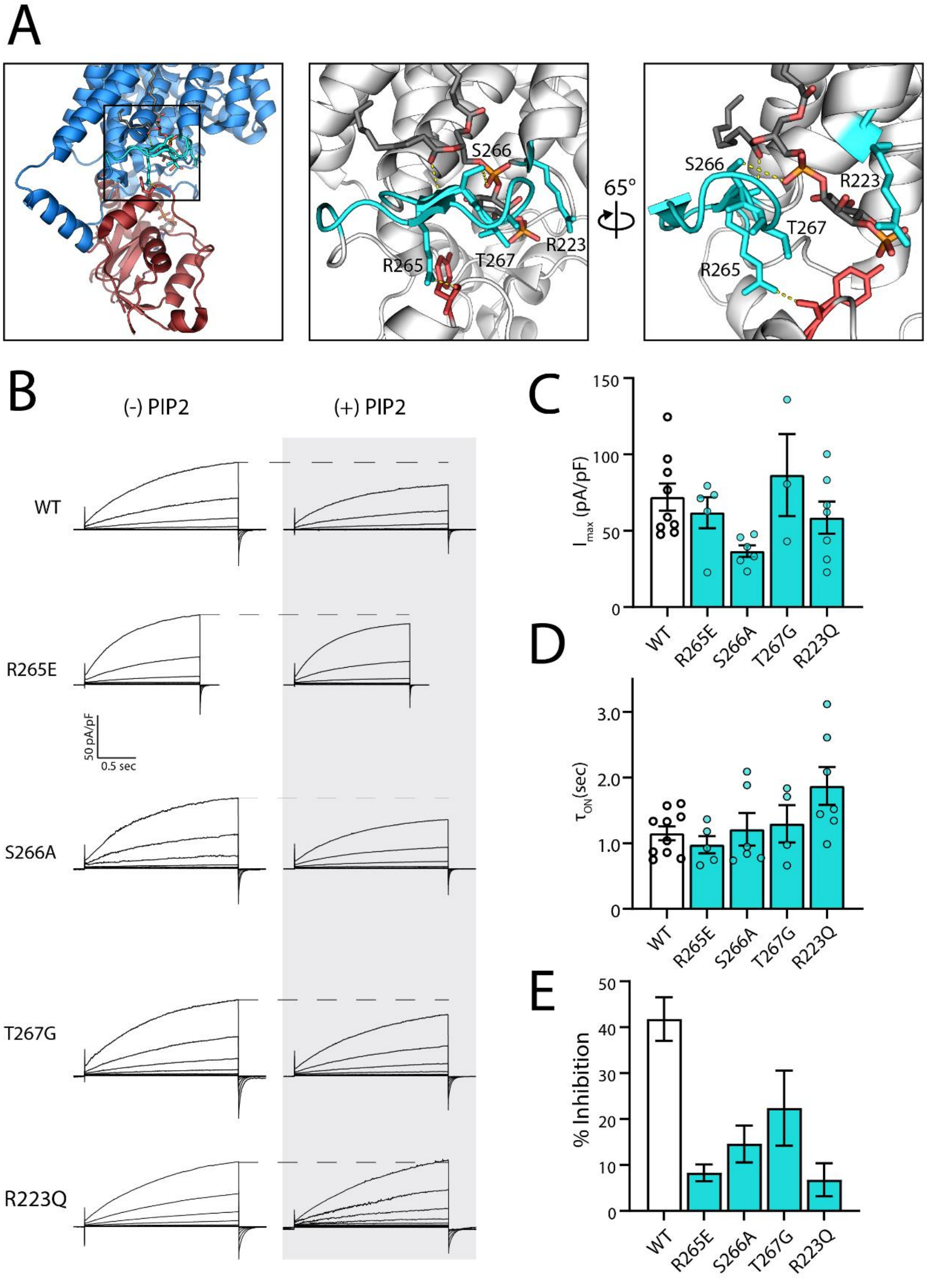


**Supplementary Figure 2. Mutations in the PIP-binding loop of ClC-7 diminish inhibition while retaining WT-like currents.** A) Views of the loop containing residues that interact with the aliphatic tail of PIP (structure 7JM7). Colors are as in Figure 1. B) Representative traces from mutants that disrupt interactions between the TMD and PIP. Mutants have diminished PI(3,5)P2 inhibition but retain characteristic slow activation and current magnitudes of WT ClC-

7. Note scale bar change compared to Figure 1. Voltage pulse protocol is as in Fig. 1C. C) Maximum currents measured after a depolarizing voltage pulse to +140 mV. Number of biological replicates is indicated in each bar and statistical significance was calculated from a Student's *t* test comparison with wild-type ClC-7 (R265E, $p$ = 0.48697; S266A, $p$ = 0.00849; T267G, $p$ = 0.50950; R223Q, $p$ = 0.34168). D) Activation time constants obtained from single exponential fits to currents at +140 mV. Replicates are indicated in each bar (R265E, $p$ = 0.33639; S266A, $p$ = 0.79881; T267G, $p$ = 0.55913; R223Q, $p$ = 0.01767). E) Percent inhibition by 50 µM PI(3,5)P2 of maximum currents measured at +140 mV, calculated as described in Fig. 1E. Biological replicates are the same as in (C).

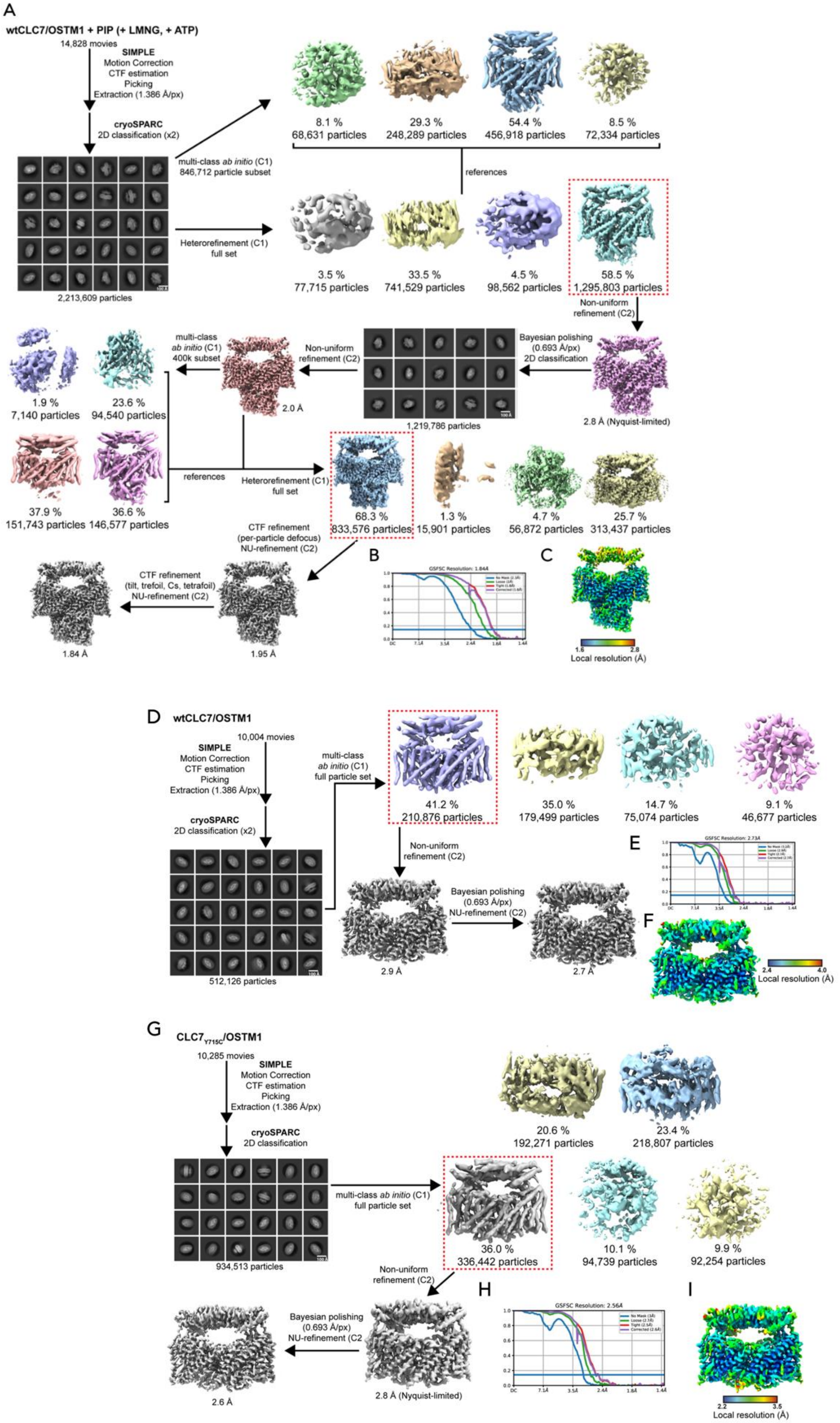
A
wtCLC7/OSTM1 + PIP (+ LMNG, + ATP)
14,828 movies
SIMPLE
Motion Correction
CTF estimation
Picking
Extraction (1.386 Å/px)
cryoSPARC
2D classification (x2)
2,213,609 particles
multi-class ab initio (C1)
846,712 particle subset
8.1 %
68,631 particles
29.3 %
248,289 particles
54.4 %
456,918 particles
8.5 %
72,334 particles
references
Heterorefinement (C1)
full set
3.5 %
77,715 particles
33.5 %
741,529 particles
4.5 %
98,562 particles
58.5 %
1,295,803 particles
Non-uniform
refinement (C2)
2.8 Å (Nyquist-limited)
Bayesian polishing
(0.693 Å/px)
2D classification
1,219,786 particles
Non-uniform
refinement (C2)
2.0 Å
multi-class
ab initio (C1)
400k subset
1.9 %
7,140 particles
23.6 %
94,540 particles
37.9 %
151,743 particles
36.6 %
146,577 particles
references
Heterorefinement (C1)
full set
68.3 %
833,576 particles
1.3 %
15,901 particles
4.7 %
56,872 particles
25.7 %
313,437 particles
CTF refinement
(per-particle defocus)
NU-refinement (C2)
1.95 Å
CTF refinement
(tilt, trefoil, Cs, tetrafoil)
NU-refinement (C2)
1.84 Å
B
GSFSC Resolution: 1.84Å
C
1.6
2.8
Local resolution (Å)
D
wtCLC7/OSTM1
10,004 movies
SIMPLE
Motion Correction
CTF estimation
Picking
Extraction (1.386 Å/px)
cryoSPARC
2D classification (x2)
512,126 particles
multi-class
ab initio (C1)
full particle set
41.2 %
210,876 particles
35.0 %
179,499 particles
14.7 %
75,074 particles
9.1 %
46,677 particles
Non-uniform
refinement (C2)
2.9 Å
Bayesian polishing
(0.693 Å/px)
NU-refinement (C2)
2.7 Å
E
GSFSC Resolution: 2.73Å
F
2.4
4.0
Local resolution (Å)
G
CLC7Y715C/OSTM1
10,285 movies
SIMPLE
Motion Correction
CTF estimation
Picking
Extraction (1.386 Å/px)
cryoSPARC
2D classification
934,513 particles
multi-class ab initio (C1)
full particle set
20.6 %
192,271 particles
23.4 %
218,807 particles
36.0 %
336,442 particles
10.1 %
94,739 particles
9.9 %
92,254 particles
Non-uniform
refinement (C2)
2.8 Å (Nyquist-limited)
Bayesian polishing
(0.693 Å/px)
NU-refinement (C2
2.6 Å
H
GSFSC Resolution: 2.56Å
I
2.2
3.5
Local resolution (Å)

**Supplementary Figure 3. Cryo-EM processing workflow of WT-CLC7/OSTM1 + PIP2, WT-CLC7/OSTM1 and CLC7-Y715C/OSTM1, including local and global resolution estimates.** (A) Image processing workflow of WT-CLC7/OSTM1 + PIP2. (C) Gold-standard Fourier Shell Correlation (FSC) curves used for global resolution estimation. (C) Local resolution estimate of the WT-CLC7/OSTM1 + PIP2 volume. (D) Image processing workflow of WT-CLC7/OSTM1. (E) Gold-standard Fourier Shell Correlation (FSC) curves used for global resolution estimation of WT-CLC7/OSTM1 volume. (F) Local resolution estimate of the WT-CLC7/OSTM1 volume. (G) Image processing workflow of CLC7-Y715C/OSTM1. (H) Gold-standard Fourier Shell Correlation (FSC) curves used for global resolution estimation of CLC7-Y715C/OSTM1 volume. (I) Local resolution estimate of the CLC7-Y715C/OSTM1 volume.

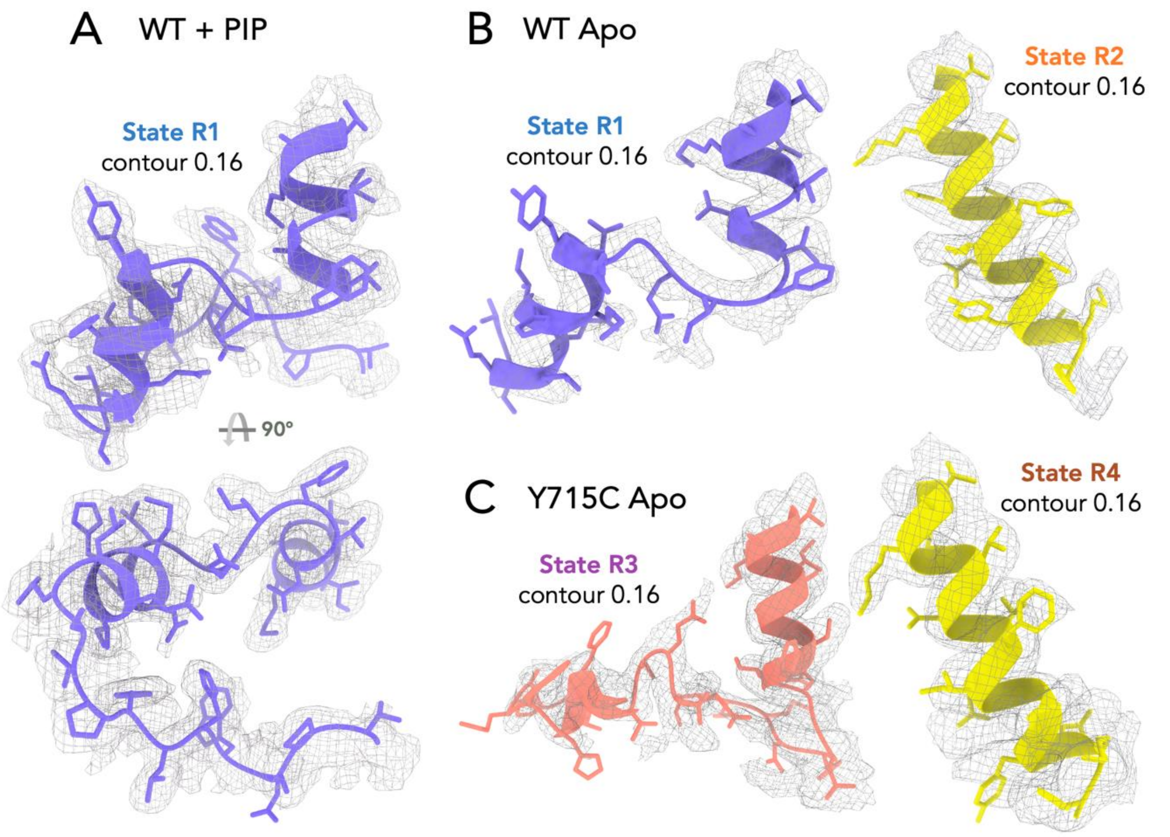


**Supplementary Figure 4. Representative densities in various Cryo EM maps of CLC7 showing the quality of chain tracing in helix Q and R.**
A. WT+PIP map; B. WT apo; C. Y715C apo.

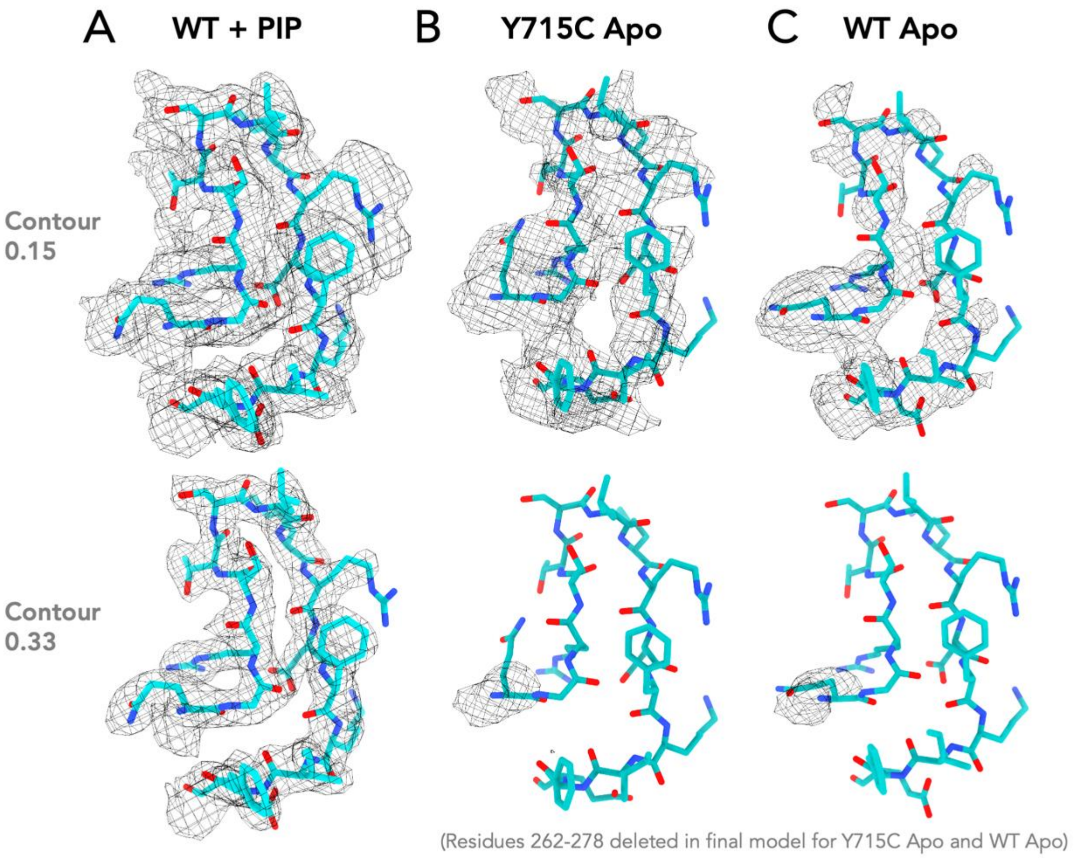


**Supplementary Figure 5. Comparison of Cryo EM densities in various CLC7 maps at F-G loop region.**

A. WT+PIP map; B. Y715C apo; C. WT apo.

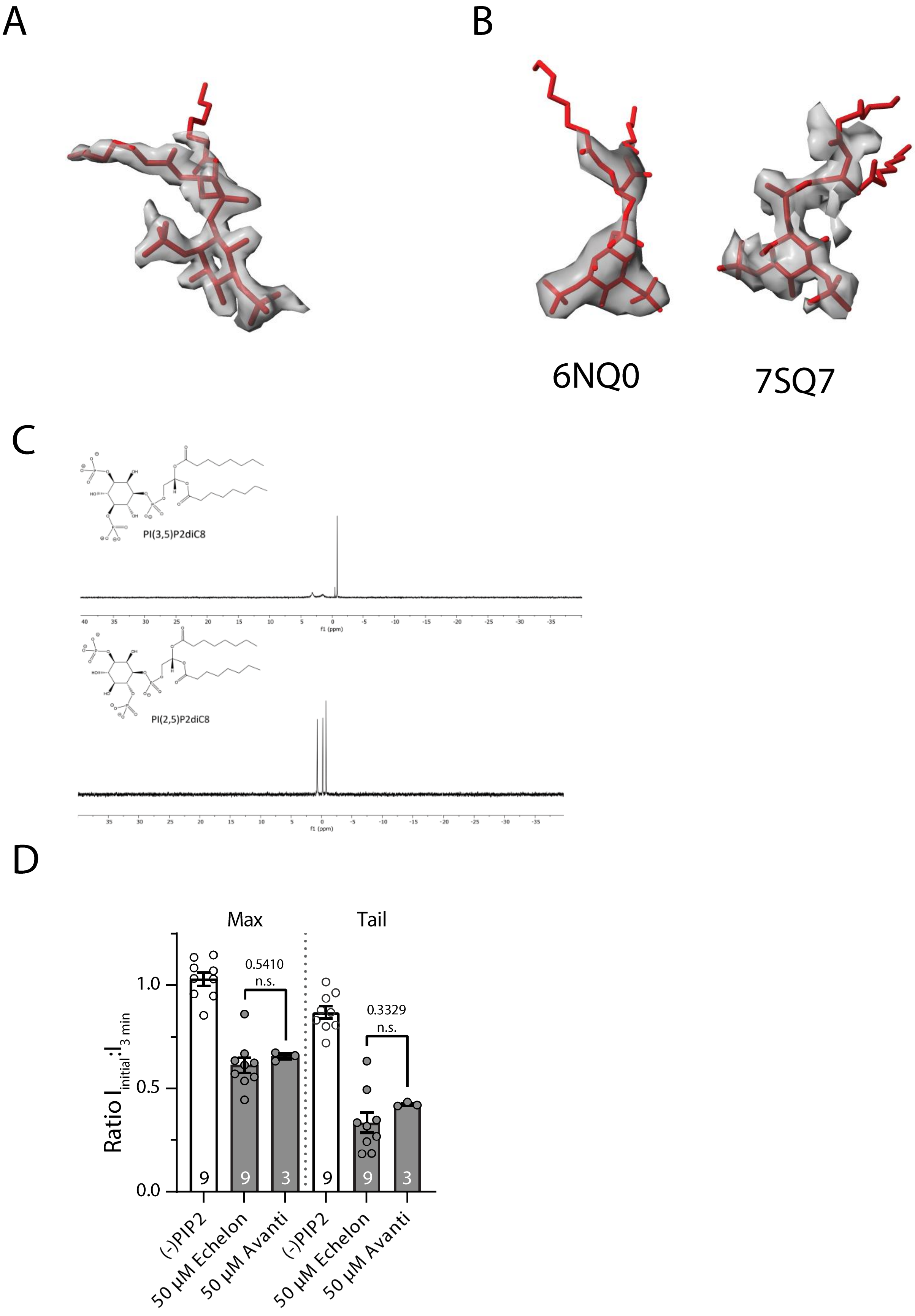
A
B
6NQ0
7SQ7
C
PI(3,5)P2diC8
f1 (ppm)
PI(2,5)P2diC8
f1 (ppm)
D
Max
Tail
0.5410
n.s.
0.3329
n.s.
Ratio I initial:I 3 min
(-)PIP2
50 μM Echelon
50 μM Avanti
(-)PIP2
50 μM Echelon
50 μM Avanti

**Supplementary Figure 6. Identification of bound lipid from the WT + PIP structure.**
A. Density of bound lipid from WT + PI(3,5)P2 structure, overlaid on a model of the lipid. Note that the a phosphate group clearly is bound to the carbon adjacent to the one carrying the acyl chains, demonstrating that this is PI(2,5)P2; this lipid was purchased from Avanti Polar Lipids. B. phosphoinositide density from two published structures (6NQ0[21] and 7SQ7[23]), with PDBs as indicated, both sourced from Echelon Biosciences. In both, the phosphates are equally spaced around the sugar chain. C. 31P NMR of lipid samples from each supplier. Top spectrum is from the Echelon sample; note the single prominent peak consistent with nearly equivalent environments for each phosphate, equally spaced around the inositol ring, consistent with PI(3,5)P2 as expected. Bottom spectrum is nominally identical PI(3,5)P2 sourced from Avanti, but note the splitting to three peaks, suggesting 3 different phosphate environments. This is incompatible with PI(3,5)P2 but is consistent with PI(2,5)P2 as suggested by the structure. D. Whole-cell patch clamp recordings compare inhibition by 50 µM PI(3,5)P2 of ClC-7PM currents by nominally PI(3,5)P2 diC8 lipids purchased from either Avanti Polar Lipids or Echelon Biosciences. Maximum currents measured at +140 mV. Values were calculated by subtracting the average percent change in current magnitude in the presence of PI(3,5)P2 from the average percent change from 30s after break-in to 3 min after break-in in the absence of the lipid and propagating the standard error of the means. Mean and Standard Error are shown, with symbols representing measurements from individual cells.

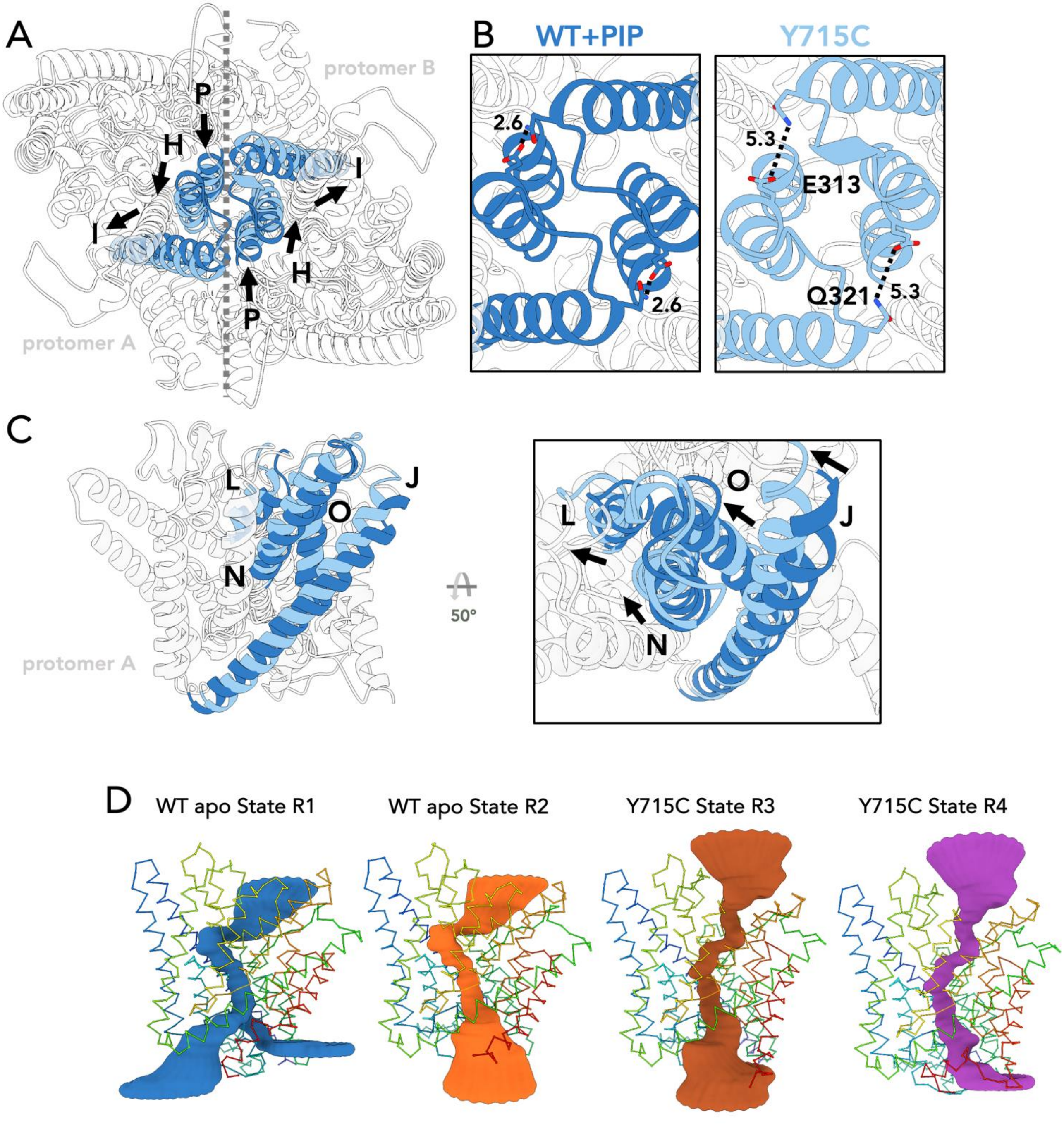


**Supplementary Figure 7. Global conformational changes between WT+PIP structure and Y715C apo.** A. CLC7 WT +PIP and Y715C apo transmembrane domain aligned, helix P, H, I coloured blue (WT+PIP: dark blue, Y715C apo: light blue) and rest of the transmembrane helices coloured white; arrows showing conformational shifts in helices. B. Close-up view of helix P, H, I, in both WT+PIP structure and Y715C apo; sidechains of residues E313 and Q321 shown and distances between these two residues across two protomers shown as dotted line. C. CLC7 WT +PIP and Y715C apo transmembrane domain aligned (only one protomer shown), helix J, L, N, O coloured blue and blue (WT+PIP: dark blue, Y715C apo: light blue) and rest of the transmembrane helices coloured white; arrows showing conformational shifts in helices.

Chloride/Proton Permeation Pathways of Different Conformational states, generated with PoreAnalyser.

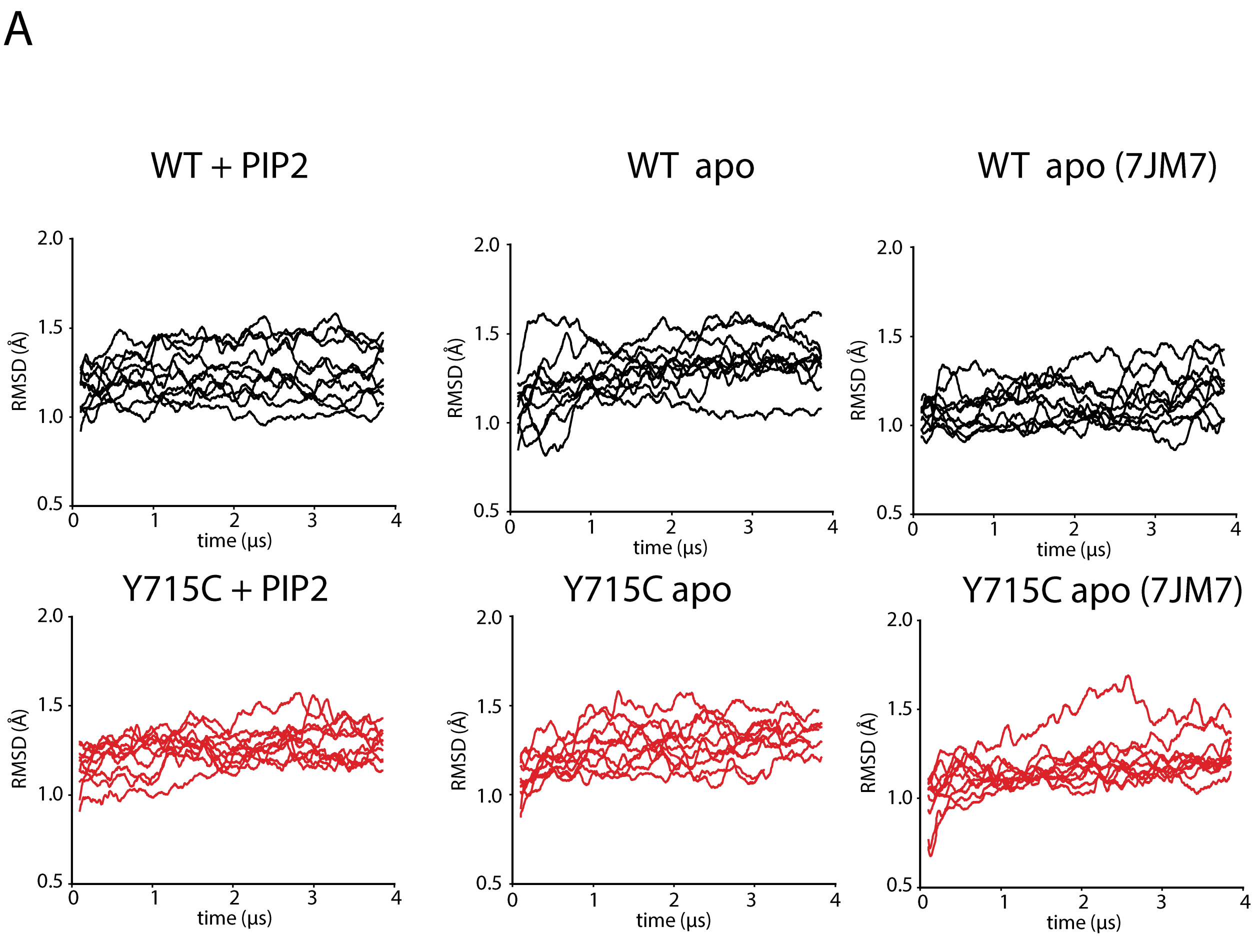


**Supplementary Figure 8. Molecular Dynamics Simulations of the ClC-7 TMD.** Root Mean Square Deviation (RMSD) time series for TMD regions from WT/WT apo/ WT apo 7JM7 (black traces) and Y715C/Y715C apo/Y715C apo 7JM7 (red traces). RMSDs were calculated relative to the TMD residues excluding loop regions of the corresponding residues in the starting WT PI(2,5)P2 cryo-EM structure. Each trace was calculated from one protomer; the 10 traces shown for each condition reflect 5 independent simulations with 2 protomers each.

**A**

| Sample | PIP | Excess LMNG:CHS | Excess ATP | CBS Domain | Resolution (Å) |
|---|---|---|---|---|---|
| wtCLC7 /OSTM1 | - | - | - | Not resolved | **2.7 (WT Apo)** |
| | + | + | + | Resolved | **1.8 (WT+PIP2)** |
| | + | + | - | Resolved | 2.8 |
| | + | - | + | Resolved | 2.8 |
| | - | + | - | Not resolved | 2.9 |
| CLC7Y715C /OSTM1 | - | - | - | Not resolved | **2.6 (Y715C Apo)** |
| | + | + | + | Not resolved | 2.8 |
| | + | - | - | Not resolved | 2.7 |

**B**

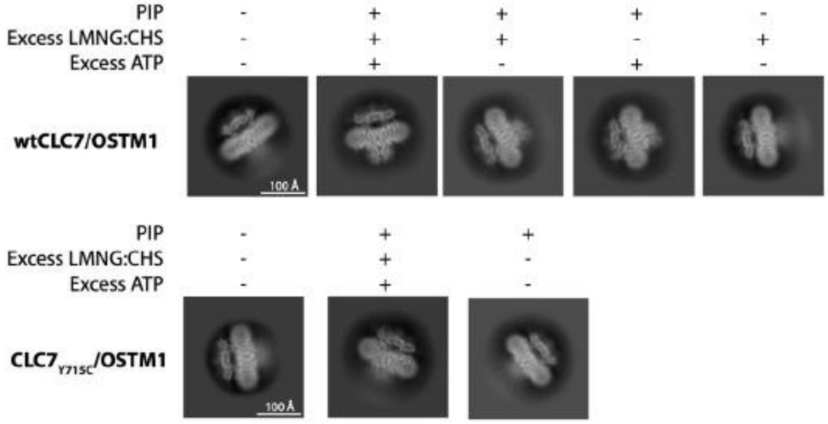


**Supplementary Figure 9. EM Datasets of WT-CLC7 and Y715C in Different Conditions.** A. Summary of the conditions (PIP2, detergent, and ATP) in different WT-CLC7 or Y715C EM datasets, the resolution of final maps, and whether CyD (CBS domain) is resolved; maps with resolution in bold were deposited. B. Examples of 2D Classes from the collected EM datasets with different conditions, showing side view of CLC7/OSTM1 complex for visualization of resolved / not resolved CyD.
